# Amyloid fibril proteomics of AD brains reveals modifiers of aggregation and toxicity

**DOI:** 10.1101/2023.03.03.530975

**Authors:** Arun Upadhyay, Deepak Chhangani, Nalini R. Rao, Julia Kofler, Robert Vassar, Diego E. Rincon-Limas, Jeffrey N. Savas

**Affiliations:** Ken and Ruth Davee Department of Neurology, Northwestern University Feinberg School of Medicine Chicago IL 60611, USA; Department of Neurology, McKnight Brain Institute, and Norman Fixel Institute for Neurological Diseases, University of Florida, Gainesville, FL, USA; Department of Pathology, Division of Neuropathology, University of Pittsburgh, Pittsburgh, PA 15213, USA; Department of Neuroscience, University of Florida, Gainesville, FL, USA; Genetics Institute, University of Florida, Gainesville, FL, USA; Mesulam Center for Cognitive Neurology and Alzheimer’s Disease, Northwestern University Feinberg School of Medicine, Chicago, IL, 60611, USA

**Keywords:** Alzheimer’s disease, amyloid, Drosophila, fibril purification, proteomics, amyloidome

## Abstract

**Background:** The accumulation of amyloid beta (Aβ) peptides in fibrils is prerequisite for Alzheimer’s disease (AD). Our understanding of the proteins that promote Aβ fibril formation and mediate neurotoxicity has been limited due to technical challenges in isolating pure amyloid fibrils from brain extracts.

**Methods:** To investigate how amyloid fibrils form and cause neurotoxicity in AD brain, we developed a robust biochemical strategy. We benchmarked the success of our purifications using electron microscopy, amyloid dyes, and a large panel of Aβ immunoassays. Tandem mass-spectrometry based proteomic analysis workflows provided quantitative measures of the amyloid fibril proteome. These methods allowed us to compare amyloid fibril composition from human AD brains, three amyloid mouse models, transgenic Aβ42 flies, and Aβ42 seeded cultured neurons.

**Results:** Amyloid fibrils are primarily composed by Aβ42 and unexpectedly harbor Aβ38 but generally lack Aβ40 peptides. Multidimensional quantitative proteomics allowed us to redefine the fibril proteome by identifying 17 new amyloid-associated proteins. Notably, we confirmed 126 previously reported plaque-associated proteins. We validated a panel of these proteins as bona fide amyloid-interacting proteins using antibodies and orthogonal proteomic analysis. One metal-binding chaperone metallothionein-3 is tightly associated with amyloid fibrils and modulates fibril formation *in vitro.* Lastly, we used a transgenic Aβ42 fly model to test if knock down or over-expression of fibril-interacting gene homologues modifies neurotoxicity. Eight RNAi lines suppressed and 11 enhanced Aβ42 toxicity.

**Conclusions:** These discoveries and subsequent confirmation indicate that fibril-associated proteins play a key role in amyloid formation and AD pathology.

## BACKGROUND

Amyloid beta (Aβ) peptides accumulate, rapidly oligomerize, and can form large degradation-resistant insoluble fibers in Alzheimer’s disease (AD) brains. Aβ peptides are generated by sequential proteolytic cleavage of the amyloid precursor protein (APP) with Aβ38, 40, and 42 being most common. Late-stage AD brains are packed with Aβ42 peptides that accumulate in a wide range of heterogeneous structures, while Aβ40 peptides are less prone to accumulate [1, 2]. The relevance of Aβ38 peptides is less clear and they may play context-dependent roles in influencing Aβ42 and Aβ40 oligomerization as well as Aβ42 toxicity [3–5]. The importance of Aβ oligomers in the etiology of AD has been established; however, we lack an understanding of how amyloid fibrils form and mature. Determining the mechanisms responsible for amyloid fibril formation may provide new and relevant insight into the development of therapeutic strategies for reducing the amyloid load.

Fibrils represent end point structural assemblies in the long process through which monomers can gradually accumulate into large aggregates and form mature plaques [6, 7]. The relevance of amyloid fibrils in AD is highlighted by the recent therapeutic success of Lecanemab, which preferentially binds to large, soluble Aβ protofibrils [8]. Thus, it’s possible that by inhibiting fibril formation or maturation, we may be able reduce the amyloid load, restore proteostasis, and even prevent neuronal death. However, the complex biochemical properties of protofibrils and fibrils (e.g., sizes, and degree of hydrophobicity), have presented a barrier to our understanding of these enigmatic proteinaceous inclusions [9]. One of the most significant barriers has been our inability to obtain highly purified amyloid fibrils from brain tissue extracts [10]. To bypass this requirement, amyloid fibril structure has been studied using synthetic Aβ peptides seeded with AD brain isolates [11, 12]. These seeding experiments produce a variety of amyloid structures but precisely how they relate to amyloid structures formed in the brain is unclear. Recently several groups have succeeded in isolating highly pure amyloid fibrils from mouse and human brains and solved their structures [13, 14]. However, an exhaustive proteomic composition of these fibrils beyond Aβ peptides has not been reported.

The formation of amyloid fibrils in the brain is a complex process that requires long time frames and culminates in the deposition of plaques predominantly near synapses in the extracellular space [6]. A wide range of proteins has been found trapped in or aggregated near plaques; however direct and indirect amyloid fibril-binding proteins are largely unknown. Previous mass spectrometry (MS)-based proteomic analyses of the Aβ interactome or the amyloid plaque proteome have reported hundreds or even thousands of proteins. Most of these studies used traditional biochemical approaches, laser microdissection, or affinity purification and captured a heterogeneous pool of amyloid-associated or coprecipitated proteins from brain, blood, or cerebrospinal fluid [15–17]. Despite these efforts, we still lack a clear understanding of the proteins involved with amyloid fibril formation and stabilization. This is primarily due to the large number of proteins and inconsistency in pools of identified proteins.

To identify proteins influencing amyloid fibril formation or modulating toxicity, we developed an amyloid fibril core purification strategy and used leading MS-based analyses to determine their content. Detailed inspection of the Aβ peptide isoforms in the amyloid fibrils revealed the presence of Aβ42, and unexpectedly Aβ38, but not Aβ40. *In vitro* studies showed Aβ38 can accelerate Aβ42 fibril formation. Inside brain, there could be other proteins present in low concentrations in the proximity of Aβ peptides influencing their aggregation and cross-reactivities. Our comprehensive proteomic analyses revealed a consistent panel of proteins associated with amyloid fibrils purified from multiple biological sources, including postmortem AD patient brains, three AD mouse models, Aβ42 overexpressing flies, and cultured neurons seeded with Aβ42 peptides. A panel of selected proteins were verified with antibodies. Among the top candidates was metallothionein-3 (Mt3), which accelerated Aβ42 aggregation *in vitro.* Finally, we confirmed that several of these proteins also regulate Aβ42-induced toxicity in a *Drosophila* model. Taken all together, our study provides a pioneering description of the amyloid fibrils and elucidates the functional influence of a panel of Aβ-interacting proteins in fibril formation and *in vivo* toxicity.

## METHODS

### Animals

Animal care and experimental protocols in this study were designed and performed as per National Institutes of Health Guidelines (protocols IS0009991). Northwestern University’s Institutional Animal Care and Use Committee (IACUC) approved these protocols. For stable ^15^N isotope labeling, previously described ^15^N isotope labeling method was followed for labeling WT animals [18]. Briefly, animals were kept on ^15^N enriched Spirulina-based diet obtained from Cambridge Isotopes Laboratories) for three months starting at P28. For euthanasia, mice were anesthetized with 3% isoflurane followed by cervical dislocation and acute decapitation. Required brain regions for each experiment were harvested, flash-frozen in a dry ice/ethanol bath, and stored at −80°C.

### Human samples

Frozen post-mortem frontal cortex tissue was obtained from the University of Pittsburgh neurodegenerative brain bank. Brain tissues were donated with consent from family members of the AD patients and approval of the University of Pittsburgh Committee for Oversight of Research and Clinical Training Involving Decedents. All institutional guidelines were followed during the collection of tissues. Staging of AD pathology was performed using NIA-AA criteria [19]. Additional details on AD patients their diagnosis, and neuropathological conditions are provided in Supplementary methods.

### Amyloid fibril purification

Biochemical purification of amyloid fibrils from mouse and human brain tissues was performed using novel technological modifications in methods described previously [20, 21]. Freshly harvested or snap-frozen brain tissues were homogenized in buffer A (0.25 M sucrose, 3 mM EDTA, 0.1 % sodium azide, and protease inhibitor cocktail in 10 mM Tris-HCl pH 7) and solubilized overnight with end-to-end rotation. For *Drosophila,* heads from flies expressing either LacZ (control) or Aβ42 using 201Y-GAL4 driver combined with nls-mcherry were snap-frozen. Before purification, fly heads were pooled into groups of sixty heads each and homogenized in buffer A. The next day, by adding dry sucrose powder, the sucrose concentration was raised to 1.2M. The solubilized tissue homogenate was then centrifuged for 45 minutes at 250,000 x g, 4°C for. After discarding the top whitish layer and intermediate aqueous layers, the pellet was dispersed in the same Buffer A with a higher 1.9 M sucrose concentration. Next centrifugation was done for 30 minutes, 125,000 x g, at 4°C. The pellet is washed twice in wash buffer (50 mM Tris-HCl) by rotating at 8,000 x g, 4°C for 15 min. Digestion buffer containing collagenase and DNase I is added to solubilize and digest the pellet for three to four hours at 37°C and washed again in the same Tris-HCl buffer. Following this, the pellet is immediately dissolved in buffer A with 1.3 M sucrose and 1% SDS. Next, solubilized pellets were centrifuged for an hour at 200,000 x g, 4°C. Pellet is saved on ice and the supernatant is centrifuged again with reduced sucrose concentration, up to 1 M, at 250,000 x g for 45 minutes. Both pellets were combined and dissolved in 200 µl Tris buffer. The aqueous solution containing highly enriched amyloid material is subjected to water bath ultrasonication in Bioruptor Pico Plus and washed five times in buffer containing 1 % SDS at 16,000 x g, 20 minutes, 4°C. The final pellet is saved and dissolved in MilliQ water or buffers per experimental requirements.

### Amyloid fibril purification from seeded primary neurons

Primary hippocampal neurons were cultured from embryonic E18 rats (Envigo). Neurons were dissociated in Papain and plated on poly-D-lysine (Sigma-Aldrich #P0899) and laminin (Gibco™ 23017015)-coated plates. Neurons were kept in Neurobasal media (Gibco™ 21103049) supplemented with SM1 (STEMCELL Technologies #05711), glutamax (Gibco™ A1286001), filtered glucose, and β-mercaptoethanol (Thermo Scientific # 0219483425) and maintained for 21 days. At DIV 21, neurons were seeded with 10 µM recombinant Aβ42 fibrils (rPeptide A-1163-2). Preformed fibrils were sonicated for ten minutes in a water bath sonicator system before seeding. Following incubation, cells were collected in the media using cell scrapers, and the above-described strategy was used to purify amyloid fibrils.

### Immunoblots

For WB, protein concentrations in each sample were measured with BCA protein Assay Kit (Thermo Scientific, Cat# 23225). Equal quantities of protein samples were boiled for five minutes in SDS Laemmli buffer. Samples were immediately loaded onto the 4-15 % Mini-PROTEAN TGX Stain-Free precast gels (BioRad # 4568084) and electrophoresed high-resolution separation of proteins based on the size. Following the electrophoresis run, the gels were used for Coomassie brilliant blue or silver staining to visualize the complete protein profile in each sample. Alternatively, transfer of total protein content onto a 0.45-micron size nitrocellulose membrane was achieved in a Bio-Rad semi-dry quick transfer apparatus. Before blocking the membranes with 5 % milk, ponceau S (Sigma Aldrich #P7170), a reversible protein binding stain, was used to observe the profile of membrane-bound proteins. After 60 minutes of blocking at RT, membranes were incubated overnight at 4°C in a required concentration of primary antibodies prepared in Tris-buffered saline with 0.1 % Tween^®^20 (TBST). The next day, following four washes in TBST, five minutes each with shaking, membranes were probed with HRP-conjugated secondary antibodies obtained from the same host. Following four TBST washes, Chemiluminescence was recorded under the Bio-Rad ChemiDoc^®^ MP Imaging system. Similarly, we performed membrane-trap dot blot analysis using a previously described method [22]. In brief, an equal amount of proteins from each sample were blotted manually on pre-activated membranes and blocked with a 5 % milk solution prepared in TBST. Antibody incubation, washing, and Chemiluminescence detection were performed similarly to WB.

### Immunostaining / immunohistochemistry

Perfusion, sectioning, and immunohistochemistry were performed as previously described [23]. Briefly, mice were transcardially perfused with PBS and drop-fixed in 4% paraformaldehyde for 24 hours. Fixed brains were then cryoprotected in 30% sucrose for at least 2 days before being embedded in Tissue-Tek OCT Compound for cryostat sectioning. Sagittal sections were prepared at 25-35 μm thickness and mounted onto gelatin-coated slides (Southern Biotech, Cat# SLD01-CS). For immunostaining, sections were kept at RT for 2 hours and then washed with PBS (3x 5 min) to remove OCT. Sections were then blocked and permeabilized with 0.2% Triton-X 100 and 10% Horse Serum (HS) in PBS for 3 hours at RT. After three PBS washes, sections were incubated overnight at 4c with primary antibodies diluted in 1% HS and 0.1% Triton-X 100. The next day, sections were washed with PBS (3 x 5 min) and then incubated with secondary antibodies in PBS. After secondary incubation, sections were washed with PBS (3x 5min) and coverslips were mounted with Fluoromount-G. Images were taken using a Nikon AXR confocal microscope at 10x and 63x.

For immunostaining of purified material, the fibrils were washed three times in 1% PBS before being blocked in 2% horse serum. After two PBS washes, fibrils were incubated overnight at 4°C with primary antibodies dissolved in PBS with 0.2% serum. The next day, fibrils were washed three times and incubated with fluorescent secondary antibodies. 10 µL of each sample were put on glass slides and observed under a Leica confocal microscope with a 63x oil objective.

### Congo red staining

Staining of fresh amyloid preparations was performed by incubating the samples with filtered 0.1% Congo red (Sigma Aldrich #C6277) solution, prepared in 50% ethyl alcohol for 20 minutes at RT. The stained samples were observed under a bright field microscope at 40X magnification.

### Amyloid kinetics experiment

For ThT-based kinetic analysis, 10 mM ThT stock solution was prepared in 1% PBS and filtered through a 0.2-micron syringe filter. Stock solutions of recombinant Aβ peptides were prepared by solubilizing lyophilized monomers (Aβ38, rPeptide A-1078-2; Aβ40:rPeptide A-1153-2; and Aβ42: rPeptide A-1163-2) in 1% NH_4_OH at 200 µM concentration as recommended by manufacturer. In a 96-well plate, the aggregation reactions (100 µL each well) were set up with 10 µM Aβ peptides and 5 µM ThT in an aggregation buffer containing 20 mM HEPES and 100mM NaCl. Additional blank wells were set up without ThT or Aβ peptides. The program in the plate reader was created to read (excitation wavelength: 440 nm, emission wavelength: 482 nm) the emission at every thirty minutes till saturation.

For seeding experiments, pre-aggregated assemblies of all three Aβ peptides were prepared by incubation in PBS for two days with end-to-end rotation at 4 °C. In 96-well plates, 10 nM seeds were added to 10 µM monomeric peptide solution. For MT3 seeding, 10 nM recombinant MT3 protein (Boster Bio Cat no. PROTP25713) was added at a in additional wells. ThT flouroscence was recorded every thirty minutes and plotted using Graphpad. Total aggregate concentrations were calculated using the secondary nucleation-dominated model in the AmyloFit online tool (https://amylofit.com) and kinetic curves were plotted in Graphpad. Values plotted on grapgh were obtained from normalization and averaging the independent values of five replicates for each reaction condition.

### ELISA assay

Multiple sandwich ELISAs were performed in ninety-six well plates provided with kits available commercially per the manufacturer’s instructions. In short, input or purified fibril samples were solubilized in 5M GuHCl for 2 hours with sonication and vortex at RT. Samples were then diluted 1:90 for NL; 1:150 for NL-F and 1:300 for NL-G-F in the standard diluent buffer. The same amount of GuHCl was added to the standard diluent buffer for blank measurements. 50 μL of blank solution, standards, and samples were loaded into antibody-coated wells and incubated for 3 hours at RT. After three washes in 1X wash buffer (provided in kits), horseradish peroxidase-conjugated antibody was added for 30 min. After three washes, the samples were incubated with stabilized chromogen for 30 min, and the reaction was stopped with an acid-based stop solution. Finally, OD was measured at 450 nm using a Synergy HTX multimode microplate reader (Biotek) and compared to a standard curve to determine the final concentration. For amyloid beta aggregated ELISA, a similar 96-well plate was prepared with 100 μL of blank, standard, and diluted test samples in a pre-coated plate with anti-aggregate antibody, which primarily captures oligomeric aggregates, but also shows reactivity for fibrils. After two hours, thoroughly washed wells were incubated for an hour with human aggregated amyloid beta biotin conjugate solution. Immediately after four washes, thirty minutes of incubation in a streptavidin-HRP working solution were done. After carefully decanting the liquid from each well washed four times, stabilized chromogen was added and stopped after thirty minutes. Finally, OD measurements for each well were taken on a microplate reader.

### Negative staining and immunogold labeling electron microscopy

For negative staining, fibrils were dissolved in MilliQ water, and 10 µL aliquot was adsorbed in duplicate on Formvar/Carbon Supported 200 mesh Copper Grids for 1-2 minutes. Following blotting, and rinsing with water, grids were immediately stained with 10 μL of 2% w/w uranyl acetate for 30 seconds. Grids were again blotted, and dried in air. Dark-field images were taken with an Eagle 4k HR 200kV CCD camera mounted on FEI Tecnai Spirit G2 transmission electron microscope (FEI) operated at 80 kV. For immunogold labeling, sample preparation was done in accordance with established protocols. Fibrils were first incubated with a blocking solution containing 0.1% Tween^®^20, 1% bovine serum albumin, 1% normal goat serum, and 0.005% sodium azide diluted in Tris-buffered saline (TBS) buffer, pH 7.4. Next, the washed fibrils were incubated with primary antibodies and control IgG antibody at 1:500 dilution for four hours at 4°C and washed thrice with PBS. Fibril-antibody conjugates were dissolved in PBS and 10 μL solution was used for adsorption on the 200 mesh Copper Grids, followed by incubation with colloidal gold secondary anti-mouse or anti-rabbit secondary antibodies for one hour. Washing with TBS and stabilization with 1% glutaraldehyde for 5 minutes were performed before counterstaining in uranyl acetate. Images were taken with FEI Tecnai Spirit G2 transmission electron microscope at 80 kV acceleration voltage.

### Proteolysis experiment

For the complete digestion of fibrils, we prepared a cocktail of multiple proteolytic enzymes with distinct specificities and wide footprints. In brief, the protease cocktail consists of α-chymotrypsin (Sigma, Cat#C3142), thermolysin (Sigma, Cat#P1512), endoproteinase Asp-N (New England Biolabs, #P8104S), Glu-C (Sigma, Cat#P2922), Arg-C (Biovendor, Cat#RBG40003005), trypsin (Promega, Cat# V5280), and Lys-C (Promega, Cat# PI90307). In a 50 µL reaction solution, 50 µg fibrils were incubated with continuous mixing with different concentrations (1X, 0.5X, and 0.25X) of protease cocktail. Concentrations of various proteases were standardized and kept in the range of 0.01 to 0.1 µg for each reaction mixture. After 30 minutes of incubation, reactions were quenched with 2X SDS buffer containing 5.2 mM PMSF & 5.2 mM EDTA, at 95 °C for 5 minutes. One-fifth by volume of each reaction mix was used for WB analysis, while the rest of the sample was reduced and alkylated before overnight incubation with trypsin for digesting remaining undigested fibril assemblies. The next day, following peptide clean-up, samples were dried and resuspended in peptide resuspension buffer to analyze 3 µg of peptides with label-free MS.

### Genetic validation in *Drosophila*

To perform functional in vivo validation of our amyloid-associated proteins from MS analysis, we utilized a well-established Drosophila model of extracellular Aβ42 deposition and toxicity [24, 25]. For this, we used a recombinant line that expresses a UAS-Aβ42 transgene in photoreceptor neurons under control of the eye-specific GMR-Gal4 driver. Thus, we crossed these recombinant Aβ42 flies with innocuous LacZ/Luciferase RNAi control transgenes and with RNAi/overexpression lines corresponding to hits from proteomics data. These crosses were cultured at 27°C throughout development, and then newly eclosed flies were observed under the microscope for phenotypic analysis in the eyes. At least five flies per genotype were randomly selected to acquire multi-focal montage imaging using Leica Z16 Apo zoom system. Transgenes that alleviate Aβ42 toxicity in the eye were categorized as suppressors, while those that make it worse were scored as enhancers. Quantification of eye phenotype was performed manually using severity scores based on eye size, depigmentation, necrosis, and ommatidial disorganization [26].

### MS sample preparation-label free quant

The protein solutions were subjected to traditional chloroform/methanol precipitation, followed by structural denaturation in 50 µL of 8 M urea dissolved in 50 mM ammonium bicarbonate (ABC) buffer. The same volume of 0.2 % ProteaseMAX (Promega, Cat# V2072) solution in ABC buffer was added and incubated for an hour with vortex. The disulfide bonds in proteins were reduced with 5 mM Tris(2-carboxyethyl)phosphine (TCEP) for 20 minutes at room temperature (RT), followed by alkylation with 10 mM iodoacetamide (IAA). Tubes were incubated in the dark for 15 minutes and immediately quenched with excess (25 mM) of TCEP prepared in ABC. Subsequently, proteins were digested overnight at 37°C using MS-grade trypsin (Promega, Cat# V5280). The next morning, digestion reaction was stopped by acidification using 1 % formic acid (FA). Desalting using C18 spin columns (Thermo Scientific, Cat# 89870) was performed per the manufacturer’s instructions. Peptide solutions were dried down in a refrigerated speed vac and stored at −80°C.

### TMT-MS Sample Preparation

We performed TMT-MS analysis following previously described methods [27]. Briefly, 100 μg of protein for each biological sample was extracted using Methanol chloroform precipitation. The protein pellets were resuspended in 6 M guanidine solution prepared in 100 mM triethylammonium bicarbonate (TEAB) buffer (Thermo Scientific, Cat# 90114). The protein solutions were reduced with 5 mM dithiothreitol (DTT) and alkylated at free SH groups of cysteine residues with 20 mM iodoacetamide (IAA). Digestion reaction for proteins was initially set up with 1 μg of MS grade LysC (Promega, Cat# PI90307) for 3 hours at RT and then continued overnight with addition of 2 μg of trypsin (Promega, Cat# V5280), at 37 °C. The next morning, the digest was acidified and desalted using C18 HyperSep columns (ThermoFisher Scientific, Cat# 60108-302). The eluted peptide solution was dried completely in a speed vac. The next day, clean peptides were resuspended in 100 mM TEAB and micro BCA peptide quantification was performed to obtain the amounts of peptides for each sample for subsequent labeling with 16 isobaric plexes of tandem mass tag (TMT) reagent. Amine reactive TMT molecules can modify the N-terminus and side chains of lysines and have been phenomenal in performing tandem mass spectrometry by multiplexing multiple samples. An equal amount of each peptide sample was incubated with individual TMT plex reagents according to the manufacturer’s instructions (ThermoFisher Scientific). After incubating for 60 minutes at RT, the reaction was stopped with 0.3% (v/v) hydroxylamine. An equal amount of isobaric labeled peptide samples were combined 1:1:1:1:1:1:1:1:1:1:1:1:1:1:1:1 and subsequently desalted with C18 HyperSep columns. The combined isobaric-labeled peptide solution was fractionated into eight fractions per manufacturer’s instructions using high pH reversed-phase peptide fractionation columns (ThermoFisher Scientific, Cat# PI84868). All eight peptide solutions were dried in a speed vac, and stored at −80°C.

### Statistical Analysis

Statistical analyses were conducted using GraphPad Prism, v9. All values in figures with error bars are presented as mean ± standard error of the mean (SEM). Comparison between groups was performed using unpaired Student’s t-tests and p-values calculated; p < 0.05 were considered statistically significant. Multiple test correction was performed with the Benjamini–Hochberg correction.

## RESULTS

### Development of a biochemical purification scheme to isolate amyloid fibrils from brain extracts

We devised our strategy by building from a previously developed sucrose-density gradient ultracentrifugation-based amyloid enrichment method to isolate pure amyloid fibrils from AD mouse models and post-mortem brain tissues (Figure 1a) [20]. To enhance extraction and improve purity of fibrils, we physically removed loosely associated proteins using ultrasonication. First, we used 5xFAD transgenic mice brains, which display a diverse collection of amyloid plaques and severe amyloid pathology. Western blot (WB) analysis with anti-fibril (LOC) and Aβ42 (Aβ_1-42_) antibodies were used to assess recovery and enrichment. Examination of the biochemical fractions across our purification revealed that the final material (i.e., P11) was highly enriched with high molecular weight (HMW) Aβ42 fibrillar species (Figure 1b and S1a). To extend our method using a more physiologically relevant model of amyloid-related pathology, we repeated these experiments using *App* knock-in (KI) mouse models containing humanized Aβ peptide amino acid sequence along with the Swedish mutation (*App^NL/NL^*), in combination with the Beyreuther/Iberian mutation (*App^NL-F/NL-F^*) and the Arctic mutation (*App^NL-G-F/NL-G-F^*) [23, 28]. To verify the presence of amyloid in these purified protein aggregates, we stained the material with Congo red (CR), an amyloid-specific diazo dye (Figure 1c). In parallel, an Aβ42 antibody confirmed that the purified amyloid fibrils were loaded with Aβ42 peptides (Figure 1d). To coarsely assess the structural diversity of the purified material, we performed negative staining electron microscopy (EM), which revealed the presence of SDS-resistant individual amyloid fibrils and fibril bundles (Figure 1e and S1b). These fibrils contained Aβ_1-42_ based on immunogold labeling (Figure 1f and S1c). We found that 5xFAD, *App^NL-F/NL-F^*, and *App^NL-G-F/NL-G-F^*, but not wild type or *App^NL/NL^*brains harbor fibrils (Figure 1g-h). Filter trap dot blot analysis with LOC (fibrils), A11 (Aβ oligomers), and 6E10 (Aβ_1-16_) antibodies also revealed the presence of amyloid fibrils (Figure S1d and S1e). Next, we quantified the insoluble Aβ peptides with solid-phase sandwich enzyme-linked immunosorbent assay (ELISA). The results indicate more Aβ aggregates in *App^NL-F/NL-F^* compared to *App^NL/NL^* brains at twelve-months, whereas *App^NL-G-F/NL-G-F^* brains had abundant aggregated deposits at both six and twelve months (Figure S1f). To test the specificity of our strategy for purifying HMW fibrillar assemblies, we isolated amyloid fibrils from *App* KI mouse brain extracts at ages with increasing degrees of amyloid pathology. Notably, a progressive deposition was observed in an age-dependent manner consistent with previous reports (Figure S1g). We extended this strategy to purify amyloid fibrils from postmortem sporadic AD human brain tissues with increasing degree of AD pathology. The specific cases analyzed were scored for AD pathology based on ABC scores [19]. A, B and C indicate amyloid spread, Braak and CERAD score, assigned between 0 and 3 (see methods for patient details). For practical reasons, patient brain samples were grouped based on Braak stages (Figure S1h). First, we quantified total amyloid aggregate loads in the homogenates of all the brains obtained (Figure S1i). WB and ELISA-based analysis revealed a similar pattern of HMW aggregates in insoluble amyloids purified from these human AD brain extracts (Figure 1i and S1j-k). Based on the results from multiple assays, we have developed a new biochemical purification strategy to isolate amyloid fibrils.

**Figure 1.**
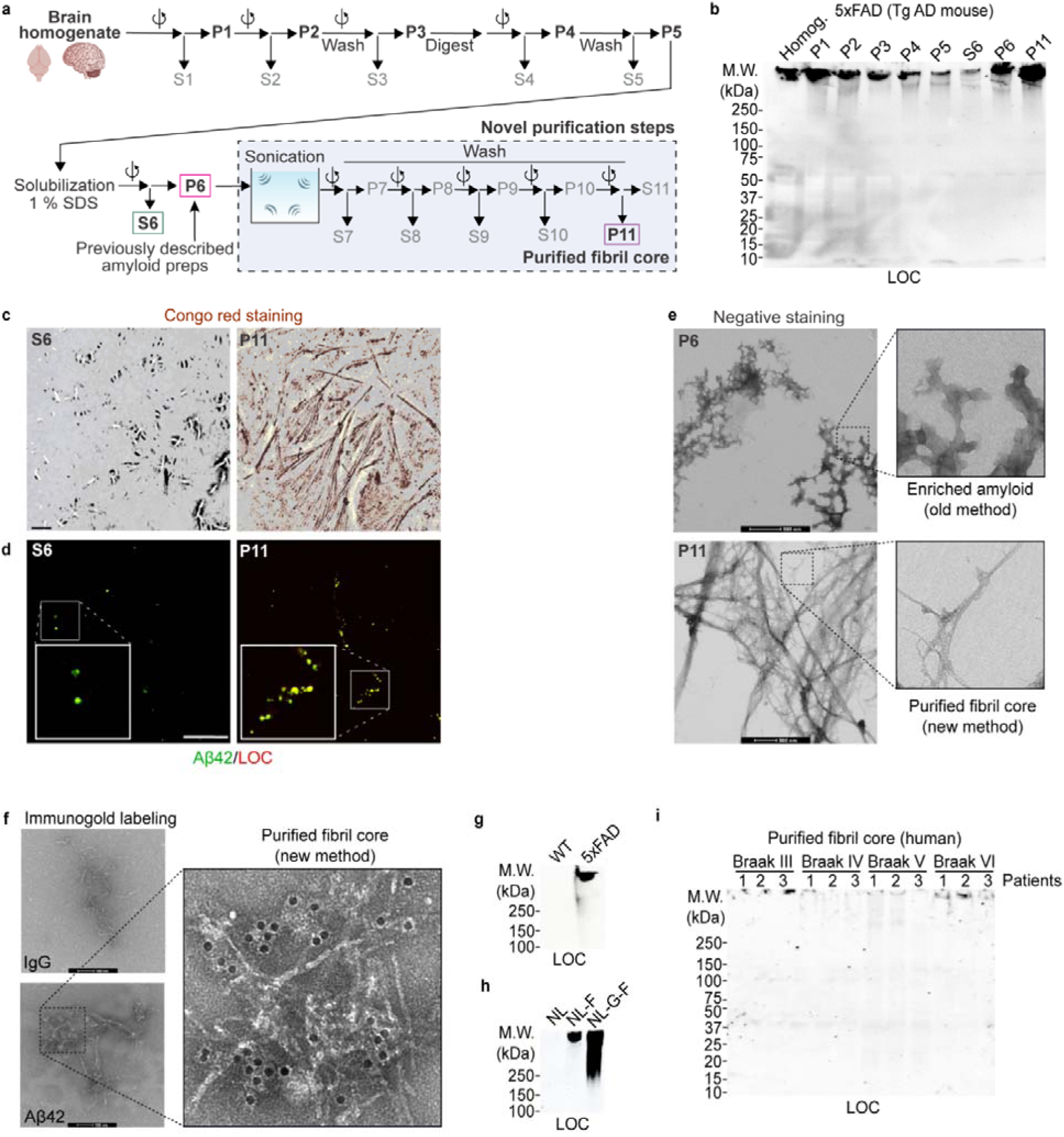
A purification strategy to isolate amyloid fibrils from AD and AD model brain extracts. (a) Biochemical purification strategy schematic. This method builds on previously developed methods and purifies SDS-insoluble dense amyloid fibril cores with sucrose density-gradient ultracentrifugation, ultrasonication, and washing with SDS. (b) Western blot analysis of indicated fractions collected during amyloid fibril purification from transgenic 5xFAD mouse cortex using anti-fibril (LOC) antibody. 10% v/v material from each fraction was loaded. (c) Amyloid specific Congo red (CR) staining of fibril cores. Bright-field images of the P11 fraction from *App^NL-G-F/NL-G-F^* mouse cortical extracts stained with amyloid-specific CR. SDS soluble (S6) fraction is used as a control. (d) Immunofluorescence (IF) images of S6 and P11 fractions using Aβ42 and LOC antibodies. (e) Representative negative staining EM analysis of amyloid material (P11) extracted using our purification strategy from *App^NL-G-F/NL-G-F^* brains compared to amyloids (P6) enriched using previously reported purification strategy. (f) Immunogold labeling with Aβ42 antibodies of purified fibrils, visualized by negative staining EM. IgG antibody was used as negative control. (g-h) WB analysis of purified fibrils isolated from WT, 5xFAD, and cortical extracts from three *App KI* (*App^NL/NL^*, *App^NL-F/NL-F^*, and *App^NL-G-F/NL-G-F^*) mouse lines with contrasting levels of amyloid pathology. (i) WB analysis of human amyloid fibrils isolated from human brain tissues with increasing pathological scores (Braak stages) with LOC antibody. Scale bar in c and d are 10 μm. P = pellet, and S = supernatant. *App^NL/NL^*= NL, *App^NL-F/NL-F^*= NL-F, *App^NL-G-F/NL-G-F^* = NL-G-F. All mice were 6 months of age.

### A**β**38 is a major component of amyloid fibrils

Aβ peptides are produced in several lengths; thus, we purified amyloid fibrils from *App^NL-G-F/NL-G-F^* brain extracts and the presence of the three most common Aβ isoforms (Aβ38, Aβ40, and Aβ42) were investigated by multiple antibody-based assays. To study the relative abundance and distribution of these three Aβ species, we collected intermediate fractions during amyloid fibril purification from *App^NL-G-F/NL-G-F^*mouse brain. First, we confirmed the specificity of all three Aβ antibodies by immunoblotting recombinant human Aβ38, Aβ40, and Aβ42 peptides, respectively (Figure S2a-b). Next, we studied the presence of individual Aβ peptide species and assemblies across the biochemical fractions (Figure 2a and S2c). Purified amyloids were highly enriched with Aβ38, and Aβ42, while Aβ40 showed least presence in fibril containing P11 fractions (Figure 2a). Based on ELISA, abundance of Aβ38 peptides were confirmed in the purified amyloid fibrils in all three AD mouse model brains (Figure 2b-e). We validated our findings in human brain amyloids across multiple Braak stages by similar assays. As evident from immunoblots, Aβ38 and Aβ42 were abundant in all Braak stages. On the other hand, Aβ40 was nearly absent in the purified amyloid fibrils (Figure 2f and S2d). Aβ ELISA analysis confirmed the immunoblots (Figure 2g-i and S2e). To further investigate the contribution of Aβ38 to amyloid fibril formation, we performed a panel of *in vitro* experiments. Through thioflavin T (ThT)-based kinetic assay and WB analysis, we examined the self-assembly and cross-seeding potential of Aβ38, Aβ40 and Aβ42 seed at very low concentrations (Figure 3a-d and S2f). This assay revealed that all three peptides, at 10 μM, self-assemble at varying rates. However, WB with LOC antibodies confirmed that only Aβ38, and Aβ42 form fibrils under these conditions. We also tested the possibility that preformed Aβ assemblies (10 nM) can cross-seed using all three Aβ peptide substrates based on ThT and WB analysis of the end-products (Figure 3e). The LOC blots confirmed that both preformed Aβ38 and Aβ42 assemblies can cross-seed Aβ42 fibril formation (see blue and pink asterisk in LOC blots). Contrary to that, Aβ40 seeds only induce formation of A11-positive oligomeric structure (green asterisk). In summary, these results indicate that amyloid fibrils are formed of Aβ38 and Aβ42 rich assemblies and at nanomolar concentration, Aβ38 can enhance Aβ42 fibril formation *in vitro*.

**Figure 2.**
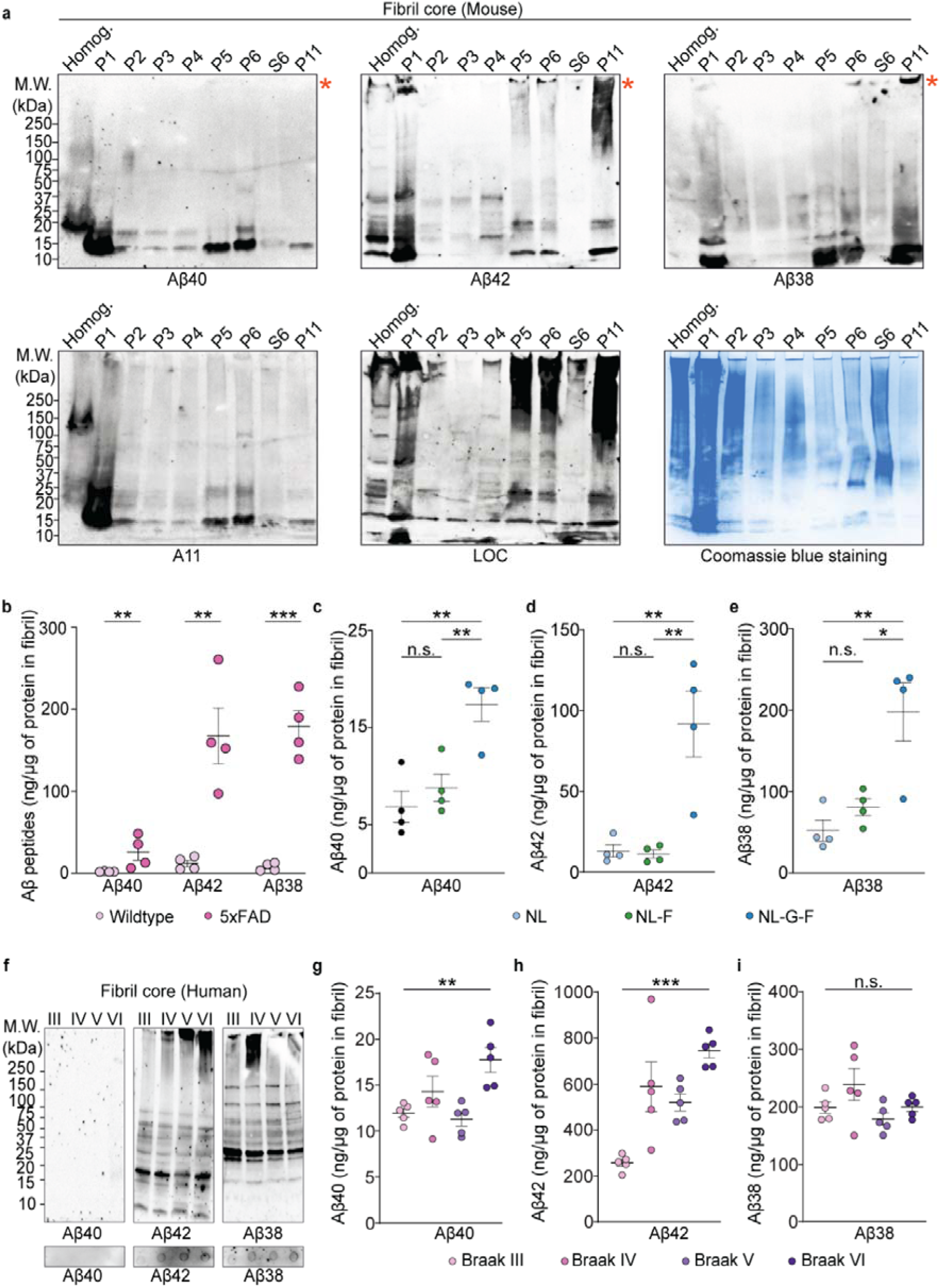
Amyloid fibril cores are enriched with Aβ42 and Aβ38 peptides. (a) WB analysis across indicated fractions from *App^NL-G-F/NL-G-F^* cortical extracts with Aβ40, Aβ42, and Aβ38 specific antibodies. A11, LOC blots, and Coomassie blue staining indicate abundance of oligomers, fibrils and total protein. Red asterisks indicate HMW aggregates. (b) Absolute quantification of Aβ40, Aβ42 and Aβ38 peptides in WT and 5xFAD purified SDS-resistant amyloid using sandwich ELISAs. (c-e) Absolute quantification of Aβ40, Aβ42, and Aβ38 peptides from *App KI* mouse brains. (f) Immunoblots of fibril cores obtained from postmortem AD brain tissues using antibodies for Aβ40, Aβ42 and Aβ38. (g-i) Absolute quantification of Aβ40, Aβ42 and Aβ38 peptides from AD human brains. Data in b-e and g-i are mean ± SEM; in B–E, n = four mice; and in G-I n= five human brain samples analyzed. *, p-value < 0.05; **, p-value< 0.01; ***, p-value < 0.001; analyzed with unpaired Student’s t-test or one-way ANOVA with post hoc Sidak test. P = pellet, and S = supernatant. *App^NL/NL^* = NL, *App^NL-F/NL-F^* = NL-F, *App^NL-G-F/NL-G-F^* = NL-G-F. All mice were 6 months of age.

**Figure 3.**
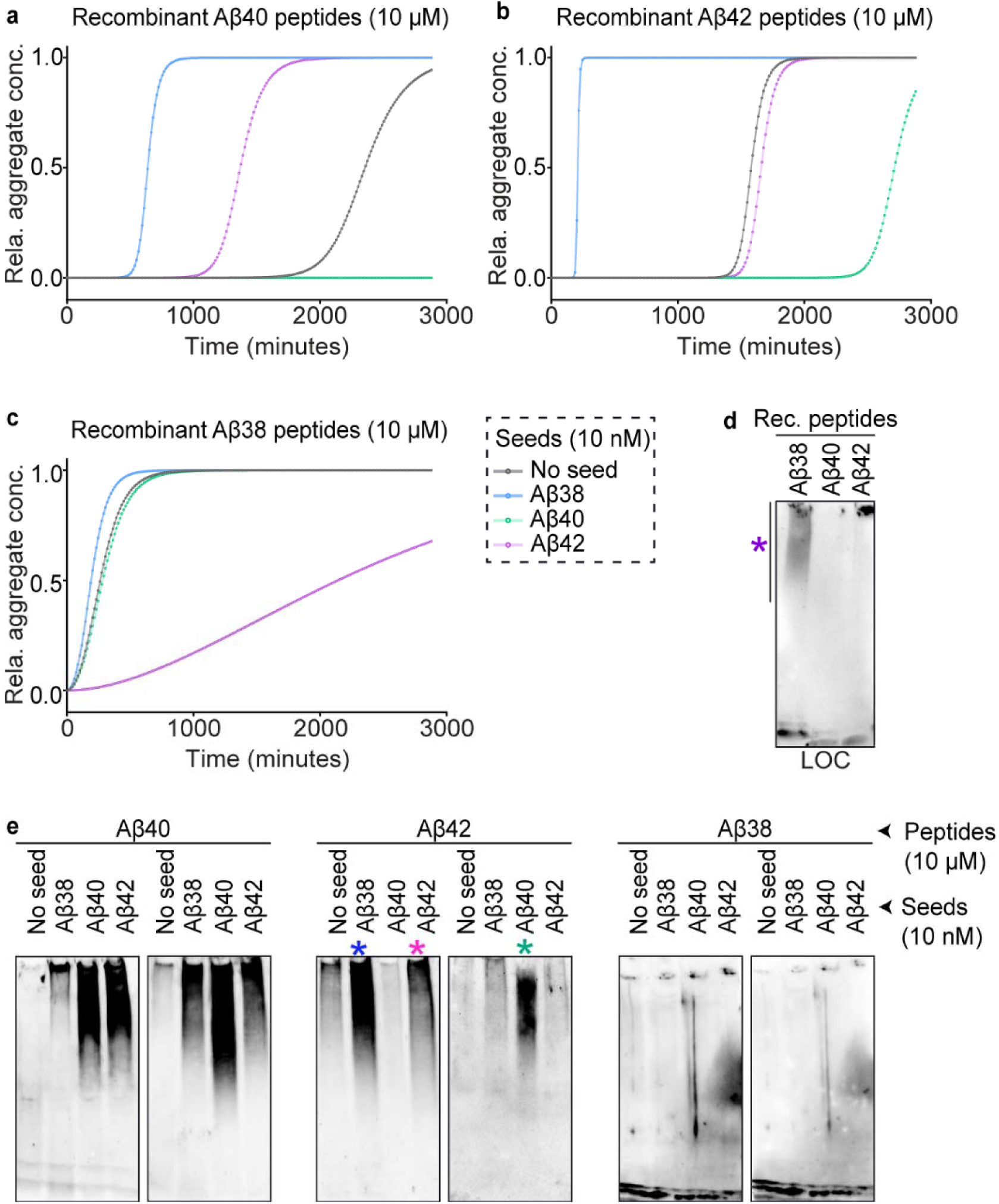
Aβ38 can seed fibril formation *in vitro*. (a-c) ThT binding kinetic assay reveals cross-seeding effects of preformed Aβ38, Aβ40, and Aβ42 fibrils on Aβ38, Aβ40, and Aβ42 substrates. Aβ38, Aβ40 and Aβ42 peptides can readily self-aggregate *in vitro* at varying rates. Additionally, Aβ38 showed self-seeding and high cross-seeding potential with Aβ42 peptides. The relative amyloid concentrations were calculated using secondary nucleation model in AmyloFit online tool (https://amylofit.com/). Five replicates were used for each sample and ThT fluorescence intensities were measured every thirty minutes. (d) WB analysis with LOC antibody reveals inherent fibril formation of recombinant Aβ38, and Aβ42, but not Aβ40, peptides *in vitro.* Aβ38 possibly forms fibrillar assemblies corresponding to lower and intermediate molecular weight (purple asterisk), while Aβ42 accumulates in highly compact high molecular weight assemblies. (e) WB analysis of end products of kinetics reaction with LOC antibody shows 10 nM seeds of Aβ42 can induce subsequent Aβ42 aggregation (pink asterisk), Aβ38 cross-seeds Aβ42 fibril formation (blue asterisk). Aβ40 accelerates formation of Aβ42 oligomers, based-on A11 antibody blots (green asterisk).

### Multiscale profiling of the amyloid fibril proteome

To identify proteins involved with the formation or stabilization of amyloid fibrils, we analyzed the purified material with MS-based proteomic analysis using five complementary workflows (Figure 4a). To confirm that our modifications to the previously reported purification strategy resulted in isolating amyloid fibrils at higher purity, we subjected the amyloid extracts like previously described methods (P6) and our purified material (P11) to label-free MS-based proteomic analysis. In all four mouse models, we significantly reduced the number of identified proteins by more than 50% compared to material prepared using previous purification method (Figure 4b). By comparing the proteins identified in the material isolated from *App^NL-F/NL-F^*, *App^NL-G-F/NL-G-F^*, and 5xFAD brains, relative to control *App^NL/NL^* brains, we delineated the proteins associated with pathological forms of amyloid and highlighted proteins identified in multiple models (Figure 4c). In the fibril cores isolated from human brain tissues, we identified the largest number of proteins from Braak stage VI, followed by Braak stage IV (Figure 4d-e). Next, we compared the relative abundance of proteins in the purified material relative to the starting material (i.e., cortical homogenate). Most proteins were identified in both analyses, with about 10% or 50% of proteins being enriched by 20-fold or more in mouse or human extracts, respectively (Figure S3a-b, Table S1-2). A panel of the most significantly enriched proteins were present in the material purified from multiple mouse models and Braak stages (Figure S3c-e). At all Braak stages, we identified a panel of peptides mapping to the APP protein and normalized the number of proteins identified relative to the number of spectra corresponding to a portion of the Aβ sequence (Figure S3f). To further ensure that the proteins identified by MS are truly associated with fibril cores rather than representing co-purifying impurities, we mixed *App^NL-G-F/NL-G-F^*brain homogenates with WT brains metabolically labeled with ^15^N spirulina chow. In this way any ^15^N protein identified must have associated in the tube and was thus deemed non-specific (Figure 4f). MS analysis revealed that > 90% of the proteins identified were ^14^N-labeled, while the remaining 10% were ^15^N-labeled (e.g., collagen, histones, titin, tubulin, myelin basic proteins and syntaxin-binding protein 1) and no longer considered as being present in the fibril cores (Figure 4g). Gene Ontology (GO) cellular component enrichment analysis revealed many of the fibril-associated proteins localize to the extracellular space (Figure 4h). To complement these studies, we also subjected the amyloid fibrils to multiple proteases to remove the proteins on the fibril periphery and to liberate peptides tightly associated with the inner fibril core (Figure S3g). We identified significantly more proteins in the 5xFAD, and *App^NL-G-F/NL-G-F^*fibrils compared to *App^NL/NL^*and *App^NL-F/NL-F^*(Figure 4i-j and Table S3). In human samples, we identified the most proteins in the purified material from Braak stage VI brains (Figure 4k-l and Table S3).

**Figure 4.**
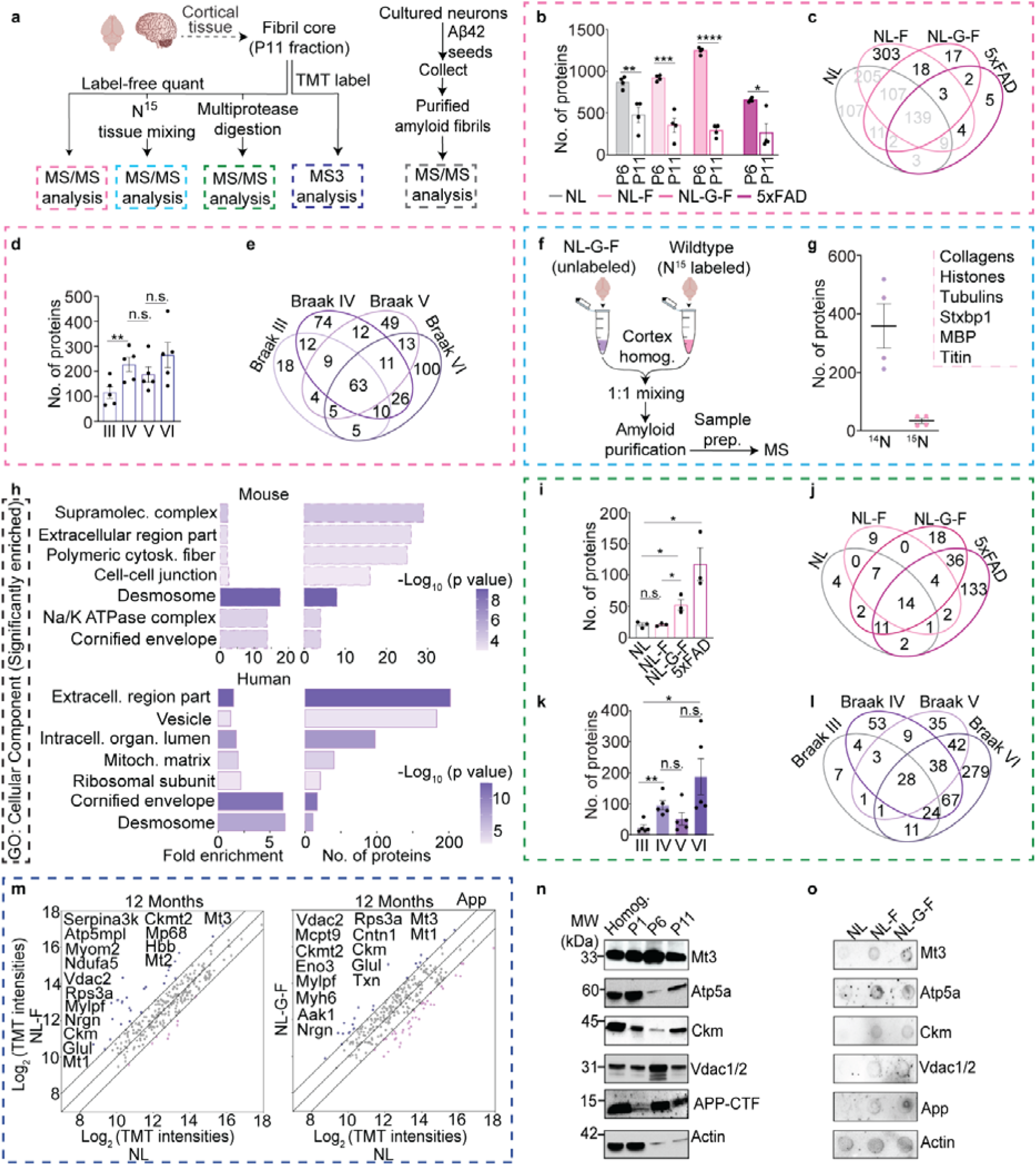
Multidimensional MS-based proteomic analyses identify amyloid fibril proteome. (a) Summary of MS analyses. (b) The new purification strategy significantly reduces the number of proteins identified in P11 fractions compared to P6 fractions collected from *App* KI (*App^NL/NL^*, *App^NL-F/NL-F^*, *App^NL-G-F/NL-G-F^*) and transgenic 5xFAD brains (n= four). (c) Venn diagram depicting the number of proteins identified in purified fibril cores across the indicated mouse strains at a 1% protein FDR. (d) Number of proteins identified in fibrils isolated from AD human brains (n= five). (e) Venn diagram comparing proteins identified in human brain amyloid fibrils. (f) Experimental workflow using ^15^N-labeled brain tissue as a control to identify nonspecific copurifying proteins. (g) Only a small panel of non-specific background proteins are identified in purified amyloid fibrils based on the identification of ^14^N (light) and ^15^N (heavy) labeled proteins; small inset shows identified ^15^N proteins. (h) The most enriched GO-cellular component associated with proteins identified exclusively in mouse and human amyloid fibrils, but not identified in input (cortex homogenates) are associated with the extracellular matrix; solid line-human, dashed line-mouse proteins. (i) Number of proteins identified in label-free MS analysis of amyloid fibril cores extracted from mouse brain tissues following digestion with multiple proteases. (j) Venn diagram comparing number of proteins identified across different mouse strains in multi-protease digestion LC-MS/MS analysis. (k) Number of proteins identified in fibril cores following multiprotease digestion of human brain-derived amyloid fibrils. (l) Venn diagram comparing number of proteins identified across human fibrils digested with multiple proteases. (m) Scatter plots comparing average TMT intensities of *App^NL-F/NL-F^* and *App^NL-G-F/NL-G-F^* with *App^NL/NL^*fibril cores. n = two biological replicates, each prepared from two pooled mouse brain cortices. (n) Immunoblots confirming the presence of selected proteins identified in the proteomic analyses. (o) Dot blots for same proteins in *App^NL/NL^*, *App^NL-F/NL-F^*, *App^NL-G-F/NL-G-F^* amyloid fibrils. Data in b, d, g, i and k represents mean ± SEM; n = three to five per analysis. *, p-value < 0.05; **, p-value< 0.01; ***, p-value < 0.001; ****, p-value < 0.0001 analyzed with unpaired Student’s t-test. P = pellet, and S = supernatant. *App^NL/NL^* = NL, *App^NL-F/NL-F^*= NL-F, *App^NL-G-F/NL-G-F^* = NL-G-F. All mice were 6 months of age unless indicated.

To obtain more rigorous quantification of the individual proteins in fibrils from all three *App* KI mouse lines at 12 and 18 months of age, we performed a 16-plex tandem mass tag (TMT) experiment (Figure S3h). We used WT (C57BL/6) cortical and *App^NL-G-F/NL-G-F^*cerebellar extracts as controls for these experiments. The biological replicates were clustered in PCA analysis based on the genotype and age (Figure S3i). Mt1, Mt3, Ckm, and Vdac2 were present at levels at least 2-fold greater in both *APP^NL-F/NL-F^* and *APP^NL-G-F/NL-G-F^* compared to *APP^NL/NL^* fibrils isolated from 12-month-old mice (Figure 4m). In fibrils from 18-month-old mice, only a putative peptidyl-prolyl cis/trans isomerase (Gm12728), a mitochondrial protein (Hadha), and the cytosolic malate dehydrogenase (Mdh1) met the same criteria (Figure S3j). Notably, Mt3 stood out as a top candidate since it was present at levels greater than 2-fold in fibrils from *APP^NL-G-F/NL-G-F^* cortex compared to the cerebellum, *APP^NL-G-F/NL-G-F^* compared to *APP^NL-F/NL-F^* at 18 months, and finally *APP^NL-G-F/NL-G-F^* at 18 months compared to 12 months (Figure S3k-m). By comparing TMT intensities of proteins identified in fibrils, we first identified proteins that were two-fold enriched in fibrils from *APP^NL-F/NL-F^* and *APP^NL-G-F/NL-G-F^* amyloids, as compared to *APP^NL/NL^*at twelve and eighteen months of age (Figure 4m, S3j and Table S4). We also homed in on proteins, which were selectively enriched in fibrils from cortex compared to those purified from the cerebellum of eighteen-month-old *APP^NL-G-F/NL-G-F^*mice (Figure S3k). Moreover, we identified proteins that were selectively enriched in fibrils from *APP^NL-F/NL-F^*with mild amyloid pathology compared to cortical fibrils from an aggressive amyloid pathology brain (*APP^NL-G-F/NL-G-F^*) of the same age (Figure S3l). Comparison of the protein levels from twelve- and eighteen-month-old *APP^NL-F/NL-F^* and *APP^NL-G-F/NL-G-F^* brains revealed proteins that bind to fibrils in an age-dependent manner (Figure S3m). Next, we validated the MS findings with a panel of antibodies and confirmed that most of these proteins are abundant in the P11 fraction (Figure 4n-o and S3n-p).

To confirm our putative amyloid fibril proteome, we incubated rodent hippocampal and cortical neurons with recombinant Aβ42 peptides and used our purification strategy to isolate amyloid fibrils and associated proteins (Figure S4a). As a first step, we performed Aβ42 WB and confirmed the presence of abundant HMW amyloid species (Figure S4b). Notably, more than two-thirds of the proteins were identified in the amyloid fibrils isolated from both cortical and hippocampal neurons (Figure S4c). Forty-nine proteins were present in both amyloid fibrils formed *in vitro* and *in vivo* (Figure S4d and Table S5). Finally, we confirmed several proteins identified in the MS and biochemistry analyses being present in amyloid plaques (Figure S4e-j). In summary, our multiscale proteomic analysis provided a short rank-ordered list of proteins physically associated with amyloid fibrils.

### Metallothionein-3 increases amyloid fibril formation

To test if the proteins we discovered closely associated with the amyloid fibril can influence fibril formation, we tested one candidate protein Mt3 that was prominent in the TMT and multiple protease proteomic datasets (Figure 4m-o and Table S3-4). Mt3 is a small cysteine-rich protein that regulates metal ions (e.g., Cu^2+^ and Zn^2+^) and is expressed primarily in the brain [29]. Mt3 levels are reduced in AD brains, but little is known about this protein’s role in amyloid pathology [30]. To confirm the presence of Mt3 in amyloid fibrils, first we confirmed an Mt3 antibody with recombinant and brain derived Mt3 proteins (Figure S5a). Following which, immunogold labeling of amyloid fibrils with the Mt3 antibody verified its presence in fibrils (Figure S5b). Next, using dot blot analysis we observed relative Mt3 protein level in *App KI* mouse brain cortex homogenates, purified fibrils and Aβ42 immunoprecipitates (Figure S5c). Immunofluorescence (IF) analysis of purified fibrils documented co-localization of Mt3 protein with Aβ42 peptides (Figure S5d). To further quantify the relative level of Mt3 in amyloid fibril cores, we performed sandwich ELISA and found the amount scaled with the amount of Aβ42 peptides in amyloid fibrils (Figure S5e-g). To investigate if Mt3 can influence Aβ aggregation, we performed *in vitro* assays with recombinant Aβ38, Aβ40, and Aβ42 peptides. Mt3 enhanced the oligomeric deposition of all three Aβ peptides and led to a reduction in LOC-positive fibril species in Mt3-containing Aβ38 and Aβ40 fibrils. No change in LOC positive fibrils was observed with Aβ42 (Figure S5h). Next, to investigate if Mt3 affects amyloid formation, we performed a ThT-based kinetic assay. We confirmed that in the presence of Mt3 protein, Aβ42 peptides aggregate at a higher rate, indicating a potential role in the formation of amyloid oligomers and protofibrils, which may act to accelerate mature fibril formation (Figure S5i). Since Mt3 is a metal chaperone and amyloid peptides associate with metal ions, we tested if the presence of metal ions (Cu^2+^ and Zn^2+^) affects binding. In this *in vitro* interaction analysis, we observed a metal-independent association of Mt3 with Aβ peptides (Figure S5j).

### Proteins associated with the amyloid fibril core modify amyloid toxicity in vivo

To assess the functional impact of the amyloid fibril-associated proteins on amyloid-induced toxicity, we used a well-established Drosophila model of Aβ42 deposition [24]. First, we tested if Aβ42 peptides form fibrils in neurons of the fly brain. For this, we collected heads from flies expressing Aβ42 in the Kenyon cells of the mushroom bodies (linked to learning and memory) using 201Y-Gal4 driver. We then purified fibrils using our newly described method (Figure 5a). WB analysis with LOC antibodies confirmed the presence of HMW amyloid fibrils in the heads of Aβ42-expressing flies (Figure 5b). MS-based proteomic analysis of isolated fibrils revealed 169 proteins with significantly higher levels compared to WT controls (Figure 5c and S6a). Additionally, 110 proteins identified in the fly amyloid fibrils are orthologs to the mammalian (25 human and 85 mouse) fibril-associated proteins (Table S6). Taken together, while not equivalent to the mammalian systems, the fly model displays similar biology in fibril formation and serves as a useful tool.

**Figure 5.**
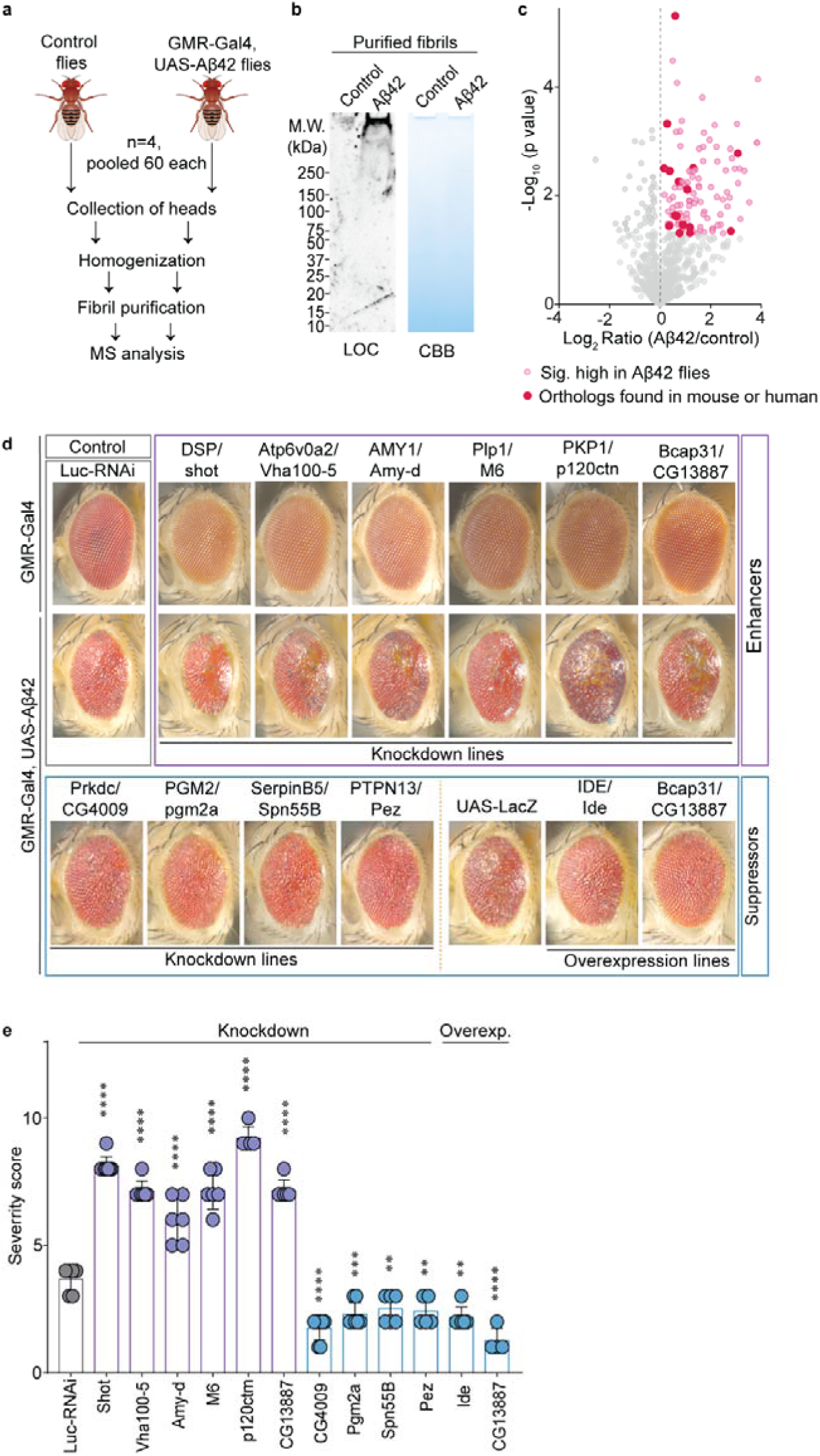
Mouse and human fibril protein orthologs interact with Aβ42 peptides *in vivo* and modulate amyloid toxicity in Drosophila. (a) Biochemical purification workflow and proteomic analysis of amyloid fibril core from transgenic flies expressing Aβ42 in adult brain using the 201Y-Gal4 driver. (b) WB analysis of the purified material isolated from LacZ control and Aβ42 flies using LOC antibody. (c) Volcano plot depicting relative abundance of proteins in fibrils isolated from flies expressing Aβ42 in adult brain compared to innocuous LacZ control flies. (d) Representative eye images of Aβ42-expressing flies carrying the indicated RNAi or overexpression lines for shortlisted genes from the MS analysis. Note that the enhancers do not modify the eye morphology in the absence of Aβ42. Luc-RNAi and UAS-LacZ were used as negative controls against RNAi and overexpression lines, respectively. (e) Histograms represent the severity scores of the indicated genetic modifiers compared to control flies. Data in e represents mean ± SD; n = five to eight representative eye images per line. **, p-value< 0.01; ***, p-value < 0.001; ****, p-value < 0.0001 analyzed with ordinary One-Way ANOVA followed by Dunnett’s multiple comparison test.

Next, we determined if modulating the expression of fly genes, orthologous to genes encoding proteins found in mouse amyloid fibrils, can modify Aβ42-induced toxicity in the fly eye. For this, we capitalized on the robust Aβ42 eye phenotype induced upon expression with the eye-specific GMR-Gal4 driver. This phenotype has 100% penetrance and is a highly reliable platform to test genetic modifiers of Aβ42-mediated toxicity [25]. Bioinformatic analysis identified 58 orthologs out of 81 mouse proteins identified in label free MS. We tested 60 RNAi or overexpression lines corresponding to those orthologs. Eight RNAi lines suppressed and eleven RNAi lines enhanced Aβ42 toxicity (Figure 5d-e, S6Bb-c and Table S6). Notably, CG4009, pgm2a, spn55b, and pez (orthologues to mouse Prkdc, Pgm2, Serpinb5, and Ptpn13 genes, respectively) showed a prominent rescue of Aβ42 insults. A small panel of fly lines over-expressing fly orthologs or human genes encoding amyloid fibril-associated proteins were obtained and crossed with GMR-Gal4>Aβ42 flies. Among these, we found two fly genes that suppress (Ide and Bcap13) and three human genes (SCAMP5, DUSP14, and LOX) that enhance Aβ42 toxicity (Figure 5d-e, S6d and Table S6).

## DISCUSSION

We set out to investigate how amyloid fibrils are formed and stabilized by mapping the amyloid fibril proteome in three model systems (mice, cultured neurons, flies) and human AD brain. Our customized biochemical purification strategy is designed specifically to isolate dense SDS-insoluble aggregates. By incorporating ultrasonication-based vibrational disruption of weakly associated proteins, combined with extensive washing, we significantly reduced the number of copurifying proteins by two-to-five-fold (Figure 4b). However, these shearing forces were not sufficiently strong to break covalent bonds or remove tightly associated binding proteins. Stable isotope labeled mouse brains served as a reliable internal standard allowing us to systematically determine the proteins that nonspecifically copurify with amyloid fibrils (Figure 4f). To validate our proteomic results, we confirmed the presence of the identified proteins by immunoblots and immunohistochemistry of three months-old *App^NL-G-F/NL-G-F^* mouse brain sections. We have thus overcome the inconsistencies previously encountered in large-scale proteomic studies and isolated fibrils with little to no contamination, which allows us to identify biologically relevant proteins, associated with amyloid fibril cores.

Previous MS-based studies have shown a sequential cleavage of APP by γ-secretase leading to generation of Aβ peptide fragments from 51 to 30 amino acids [31]. The shorter peptides (e.g., Aβ38 or Aβ40) are historically considered less-toxic and unable to cause behavioral deficits in fly and mouse models [5]. In fact, application of γ-secretase modulators (GSMs) leads to a decrease in larger Aβ peptides that are both substrates and products of γ-secretase [32, 33]. Considering targeting γ-secretase enzyme as an anti-AD therapeutic strategy, several attempts have been made in the past and are still ongoing with some success [34, 35]. However, other evidence indicates production of Aβ38 in Aβ42-independent manner, contradicting precursor-product relationship among the two and abolishing effects of multiple GSMs [36, 37]. Importantly, Aβ38 peptides have been detected at extracellular amyloid plaques in both sporadic and familial AD patients and mouse models [38]. Additionally, recent MS-based imaging of *App^NL-G-F/NL-G-F^*brains showed that Aβ38 is deposited specifically during plaque growth and may crosstalk with Aβ42 peptides [39]. Notably, shorter Aβ peptides (Aβ37, Aβ38, and Aβ40) can modulate Aβ42 fibril formation at high concentrations [40]. However, in patients the situation is highly complex since a lower risk of AD-related changes was observed in patients with high CSF Aβ38 levels [41]. In our studies, we found Aβ38 peptides are highly abundant in amyloid fibrils purified from mouse and AD human brains. More importantly, the bulk level of Aβ38 peptides corelated to a higher degree with total Aβ aggregates compared to Aβ40 or Aβ42 (Figure S1i, S2e). Somewhat unexpectedly, a cross seeding effect of Aβ38 on Aβ42 peptides was also observed. These three observations provide potential insight into APP processing and Aβ aggregation dynamics.

Most *in vitro* studies performed at micromolar concentration and show a prompt aggregation of Aβ peptides; however, brain harbors these peptides only in nanomolar concentrations. That’s why the peptides take years to decades to deposit and form long fibrils. We hypothesize that there could possibly be other cellular proteins in the proximity of Aβ peptides that assist in the initial oligomerization and nucleation. We performed a line of experiments to comprehensively investigate the core proteome of the highly pure amyloid fibrils. Overall, one hundred forty-three proteins reproducibly identified in Aβ fibrils (from two or more sources) provide a unique perspective on where and how amyloid fibrils are formed and cause toxicity. Notably, one hundred twenty-six of these proteins have previously been found to co-purify or co-localize with Aβ42 in the brain, indicating consistency between our results and several previous studies. Biochemical evidence confirming a direct physical interaction with Aβ42 peptides is lacking for most proteins (Table 1). Many of these proteins localize to the extracellular matrix, cellular membranes, and axon terminals (Figure S7), which is in line with several previous reports aimed at studying amyloid coronae [42]. The highly abundant proteins spectrin, actin, and dynein have all been previously found to be associated with amyloid plaques using antibody-staining [43, 44]. Similarly, a neurofilament protein alpha-internexin and NCK-interacting protein with SH3 domain (NCKIPSD), an actin-binding proteins are deposited in Aβ-positive dystrophic neurites [45, 46].

**Table 1:**
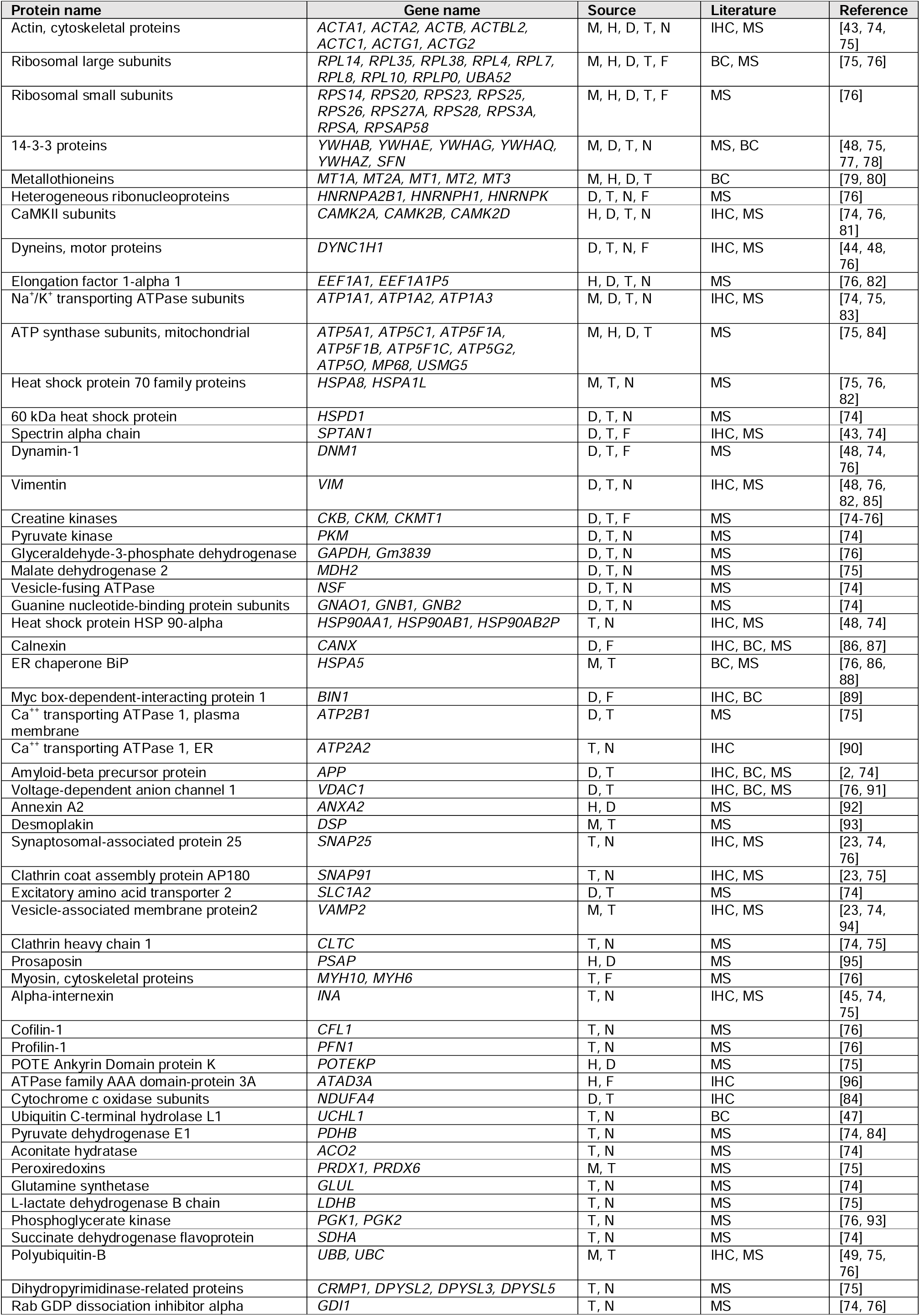
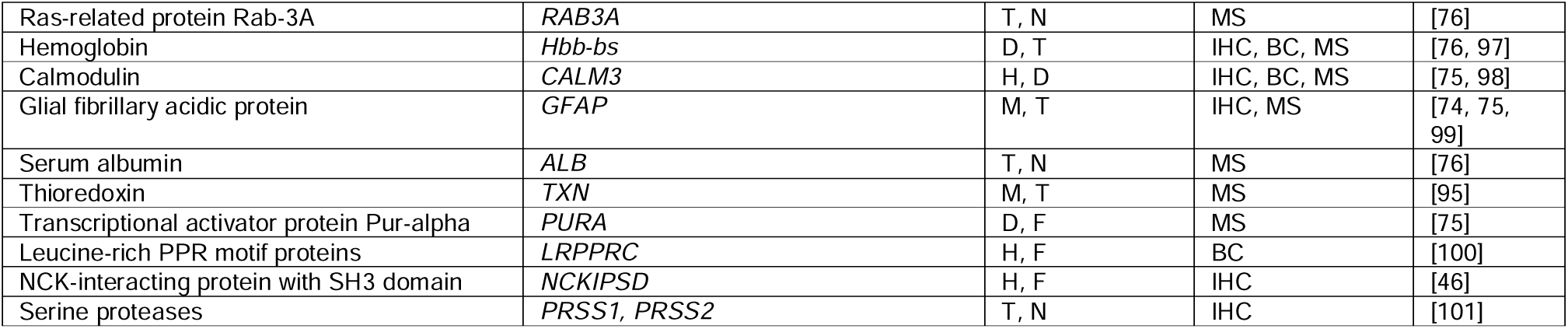
Summary of amyloid fibril associated proteins that have been previously found at amyloid plaques or with Aβ peptides. Proteins reproducibly identified in multiple proteomic analyses of amyloid fibrils isolated from brains (mouse, human, and Drosophila) or rat neurons using label-free or TMT 16-plex MS3-based quantification. Experimental evidence in this study-M: Mouse (LFQ MS/MS); H: Human (LFQ MS/MS); D: Multiprotease digestion (LFQ MS/MS); T: TMT analysis (MS3); N: primary neurons (LFQ MS/MS); F: Fly (LFQ MS/MS). Available literature evidence-MS: proteomic study; IHC: Immunohistochemistry; Biochem: Biochemical interaction.

Consistent with previous reports, we found many proteostasis-related proteins associated with amyloid fibrils, including HSP70, HSP60, HSP90 chaperones, and ubiquitin proteasome system components, such as ubiquitin and UCHL1 [47–49]. We speculate that these proteins interact with Aβ peptides soon after they start misfolding or accumulating, probably inside of the cells to circumvent the proteotoxicity. The presence of these and other intracellular proteins is at odds with extracellular amyloid deposition but consistent with previous findings on the intracellular production of Aβ peptides [50–52]. It is also possible that these molecular chaperones were extracellularly exported through non-conventional secretory mechanisms [53]. Several proteins associated with the endoplasmic reticulum (ER) (e.g., calnexin and ER chaperone BiP / Hspa5) were also well represented in fibril proteome datasets. The ER is a possible location for APP cleavage and generation of Aβ peptides in neurons, further supporting the possibility that oligomers and amyloid fibrils start assembling with in neuronal cells [50, 54]. The calcium/calmodulin-dependent protein kinase II (CaMk2a), which is a known kinase responsible for APP phosphorylation, was also found in the fibrils [52, 55]. In our analysis, we found that overexpression of insulin degrading enzyme (IDE) and knockdown of a protease inhibitor Serpinb5 both could suppress Aβ-induced toxicity in Drosophila eye neurons. Interestingly, we found that siRNA gene knock down of Bcap31 (an ER transmembrane protein) enhanced toxicity while overexpression rescued toxicity. Consistently, knock out of Bcap31 in APP / PS1 transgenic mice increased the Aβ plaque load [56]. Metallothioneins are low molecular weight (LMW) cysteine-rich metal-stabilizing proteins that have been implicated in a wide range of functions in diverse tissues. However, the functionally distinct, brain-specific isoform Mt3 is a small 68 amino acid-long metal-binding chaperone protein that was initially discovered as a neuroinhibitory factor [29]. We confirmed its presence in fibril cores, while the levels in brain homogenates were almost undetectable in our dot blot analysis. Recombinant Mt3 protein accelerated oligomer formation in all three Aβ peptides tested, while no effect on fibril formation tendency was noted. This effect does not seem to depend on metal ions.

We identified seventeen proteins that have never been found in amyloid fibrils or plaques (Table 2). Notably, three of these proteins (i.e., endophilin-A1, amphiphysin, and calcium-dependent secretion activator 1) are concentrated at axon terminals and are associated with the synaptic vesicle cycle [57–59]. Overexpression of endophilin A1 leads to Aβ accumulation, synaptic alterations and cognitive decline, while reducing ACAT1 can inhibit Aβ production in AD mouse models [60, 61]. On the other hand, peptidyl-prolyl cis-trans isomerase A (PPIA), a blood brain barrier regulatory protein, is protective effects against Aβ-induced toxicity [62]. The mitochondrial chaperone Hspe1 and TCA enzymes, 6-phosphogluconate dehydrogenase (PGD) and isocitrate dehydrogenase (IDH3B) have altered expression in postmortem AD subjects compared to healthy controls [63–65]. AD patient brain capillaries and blood respectively have elevated levels of ER translocon protein SSR4 and Y chromosome linked protein DDX3Y [66, 67]. Similarly, CLIP-associating protein 1 (CLASP1) and cAMP-dependent protein kinase II-β regulatory subunit (PRKAR2B) gene expression is elevated in APP/PS1 mouse brains [68]. In addition, we found the ADP ribosylation factor ARF5 (ER trafficking GTPases) associated with amyloid fibrils. ARF5 has never before been implicated in AD; however, ARF6, a paralog, plays an important role in APP cleavage by affecting BACE1 endosomal sorting [69]. Bifunctional purine biosynthesis protein ATIC mutation causes differential expression of several AD related genes [70]. Splicing factor SRSF4, and pseudogene HMGB1P1 are associated with frontotemporal dementia and Huntington’s disease and may have potential roles in AD pathology through tau [71, 72]. These observations prove that our findings are relevant to AD etiology and pathology.

**Table 2:**
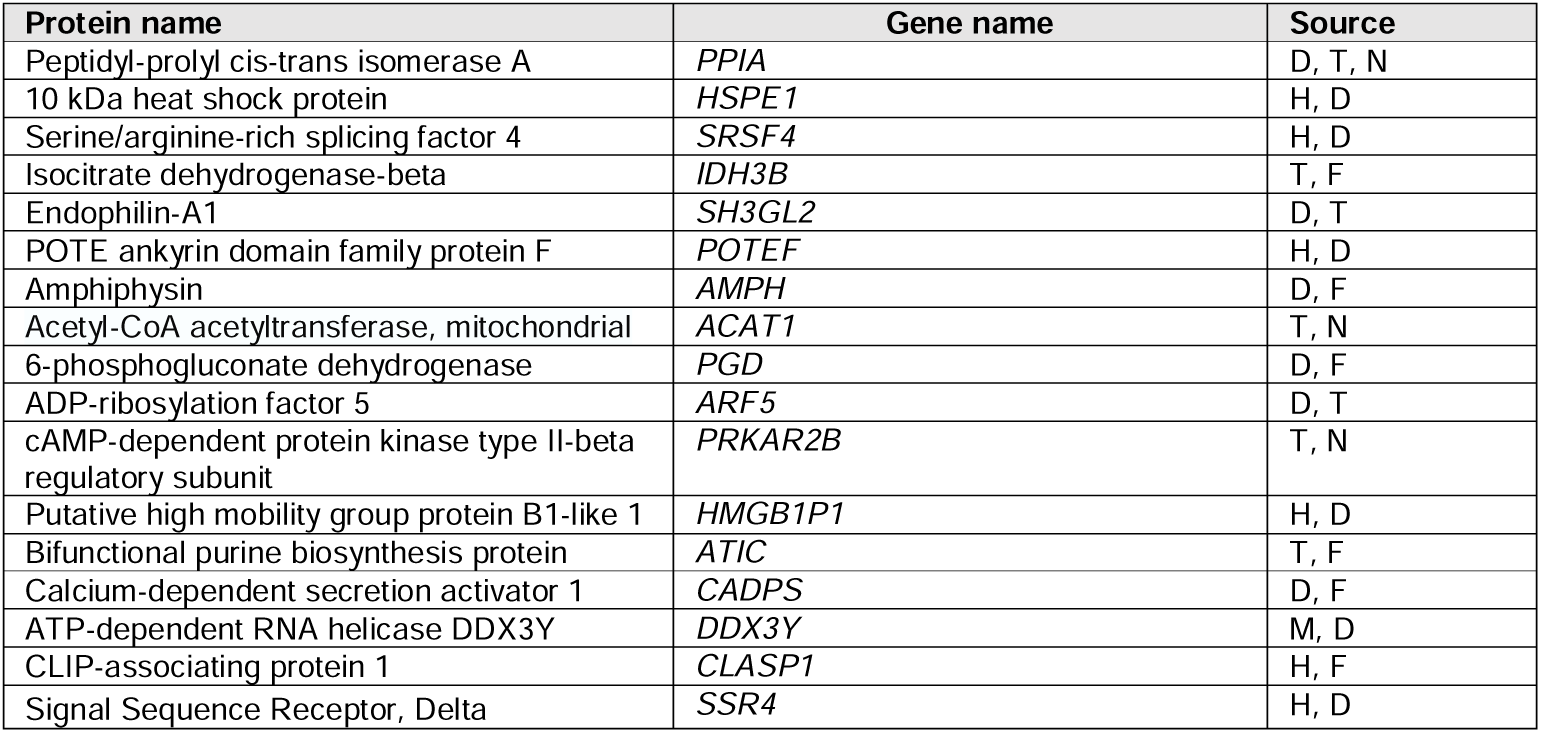
Newly discovered proteins found associated with purified Aβ fibrils. Summary of proteins identified in our analyses that have not yet been reported to be associated with Aβ fibrils. Experimental evidence in this study-M: Mouse (LFQ MS/MS); H: Human (LFQ MS/MS); D: Multiprotease digestion (LFQ MS/MS); T: TMT analysis (MS3); N: primary neurons (LFQ MS/MS); F: Fly (LFQ MS/MS).

We hypothesized that by identifying proteins tightly associated with amyloid fibrils, we would be able to strengthen our understanding of how these pernicious structures are formed, cause neurotoxicity, and may be targeted for therapeutic benefit. The consequence of these proteins associating with Aβ peptides and structural assemblies causes a loss-of-function effect by reducing the pool of functional proteins. Notably, we identified the insulin degradation enzyme (Ide), which degrades Aβ peptides and may represent one such example [73]. On the other hand, some proteins may exhibit a gain-of-function effect when their early interaction with Aβ peptides may alter amyloid plaque formation. For example, the protein quality control machinery is likely to have a significant effect on the initiation, maturation, or stabilization of nascent amyloid seeds. However, we acknowledge that some amyloid-associated proteins likely accumulate over time in the large amyloid deposits/plaques and may not have active participation in amyloid formation, elongation, or maintenance. Such interactions may be attributed to the high hydrophobicity generated in the amyloid nanoenvironment because of the presence of fibrils/plaques in the vicinity.

In summary, ultrasonication proved to be a robust agitating strategy to remove proteins weakly associated with amyloid fibrils and allowed us to comprehensively study the amyloid fibril proteome. We discovered Aβ38 in significant abundance in Aβ42-laden fibril cores, while the highly studied Aβ40 was mostly absent. While most previous reports have found that amyloid forms in the extracellular space, intracellular formation has also been found to play a key role [50, 51]. Our analysis establishes interaction between Aβ42 peptides and other proteins that are tentatively considered intracellular. Multiple experiments identified MTs in amyloid fibrils. Mt3 protein was also found to be effective in triggering deposition of Aβ42 aggregates. We postulate a similar interaction pattern of other proteins identified in this analysis with Aβ42. Some of these interactions may contribute towards stability and longevity of the fibrils. Finally, we confirmed the *in vivo* adequacy of some of the identified proteins towards targeting Aβ42-associated toxicity in a relevant AD fly model. The genetic association was established by RNAi lines when some of these showed a more aggressive phenotype when co-expressed with Aβ42 in the Drosophila eye. Remarkably, a handful of knockdown lines presented a rescue effect in the form of reduced toxicity. Future studies may help delineate more such proteins and identify modulators of Aβ42 aggregation and toxicity. We believe this work provides a foundation for more studies to identify close interaction partners and effective modulators of Aβ42 aggregation. Targeting these proteins may provide highly effective therapeutic tools to develop new AD treatments.

## CONCLUSIONS

Our novel biochemical amyloid purification strategy reduced the number of co-purifying non-specific proteins by up to five-fold. Biochemical assays confirmed presence of Aβ38 in fibrils isolated from brain and that Aβ38 can influence Aβ42 fibrilization *in vitro*. A comprehensive proteomic analysis identified 143 high confidence proteins that interact with Aβ42 during early deposition or formation of amyloid fibril cores. Most importantly, we identified 17 Aβ42-interacting proteins, which have never previously been reported in amyloid plaques. To test if the newly discovered fibril associated proteins play a functional role we followed up on the metal-binding protein Mt3. Interestingly, this protein, apart from showing high abundance in amyloid fibrils, promoted Aβ42 fibrilization *in vitro* in a metal-independent manner. Notably, knockdown of the Bcap31 fly homologue aggravated while overexpression rescued Aβ42-induced toxicity in *Drosophila* eye neurons. Similarly, overexpression of IDE (a protease) and knockdown of Serpinb5 (a protease inhibitor) also rescue toxicity in the Drosophila model. Overall, the results from our study identified several novel Aβ42-associated proteins that modify Aβ42 amyloid formation and influence neurotoxicity.

## LIST OF ABBREVIATIONS

Aβ: Amyloid beta
AD: Alzheimer’s disease
ARF5: ADP ribosylation factor 5
Bcap31: B-cell receptor-associated protein 31
CERAD: Consortium to establish a registry for Alzheimer’s disease
CLASP1: CLIP-associating protein 1
CR: Congo red
DDX3Y: DEAD box protein 3, Y-chromosomal
ELISA: Enzyme-linked immunoassay
GSM: Gamma-secretase modulators
HMGB1P1: High-mobility group box 1 pseudogene 1
HMW: High molecular weight
Hsp70: Heat shock protein 70
IDE: Insulin degradation enzyme
IDH3B: Isocitrate dehydrogenase
LC-MS: Liquid chromatography-mass spectrometry
LFQ: Label-free quantification
LMW: Low molecular weight
Mt3: Metallothionein
NCKIPSD: NCK-interacting protein with SH3 domain
PGD: 6-phosphogluconate dehydrogenase
PPIA: Peptidyl-prolyl cis-trans isomerase A
SRSF4: Serine and arginine rich splicing factor 4
ThT: Thioflavin T
TMT: Tandem mass tag
Uchl1: Ubiquitin C-terminal hydrolase L1

## DECLARATIONS

### Ethical approval and consent to participate

All the mouse and *Drosophila* work was approved by the ethics committee of Northwestern University and University of Florida, respectively. Brain tissues were collected with consent from family members of the AD patients and approval of the University of Pittsburgh Committee for Oversight of Research and Clinical Training Involving Decedents. All institutional guidelines were followed during the collection of tissues.

### Consent for publication

All authors have given their consent for publications.

### Availability of data and materials

All data are available in the main text or the supplementary information files. Experimental procedures, methods of data collection and analysis are provided in Additional file 2. The analyzed MS datasets for individual MS experiments are provided in supplementary Tables S1-S6. All raw mass spectrometry data will be released on MassIVE and Proteome Exchange following acceptance of the article.

### Author contributions

AU and JNS conceived and design of this study. AU, DC, NR and JK acquired the primary data. AU, DC, DER, RV and JNS analyzed the primary data. AU and JNS wrote the original draft of the manuscript and all authors contributed and substantively revised the manuscript. All authors read and approved the final manuscript.

## Acknowledgements

We sincerely thank Drs. Ansgar Siemer and Ralf Langen for their crucial input during development of the purification protocol. We thank Akhil Patel for technical assistance in the fly experiments and Dr. Farida Korobova for help with imaging at Northwestern University Center for Advanced Microscopy. Authors thank Vassar and Savas research group members at Northwestern University for thoughtful discussions.

