## Additional File 1 for "Amyloid fibril proteomics of AD brains reveals modifiers of aggregation and toxicity"

**SUPPLEMENTARY INFORMATION FILE 1**

**Supplementary Figures and Tables**

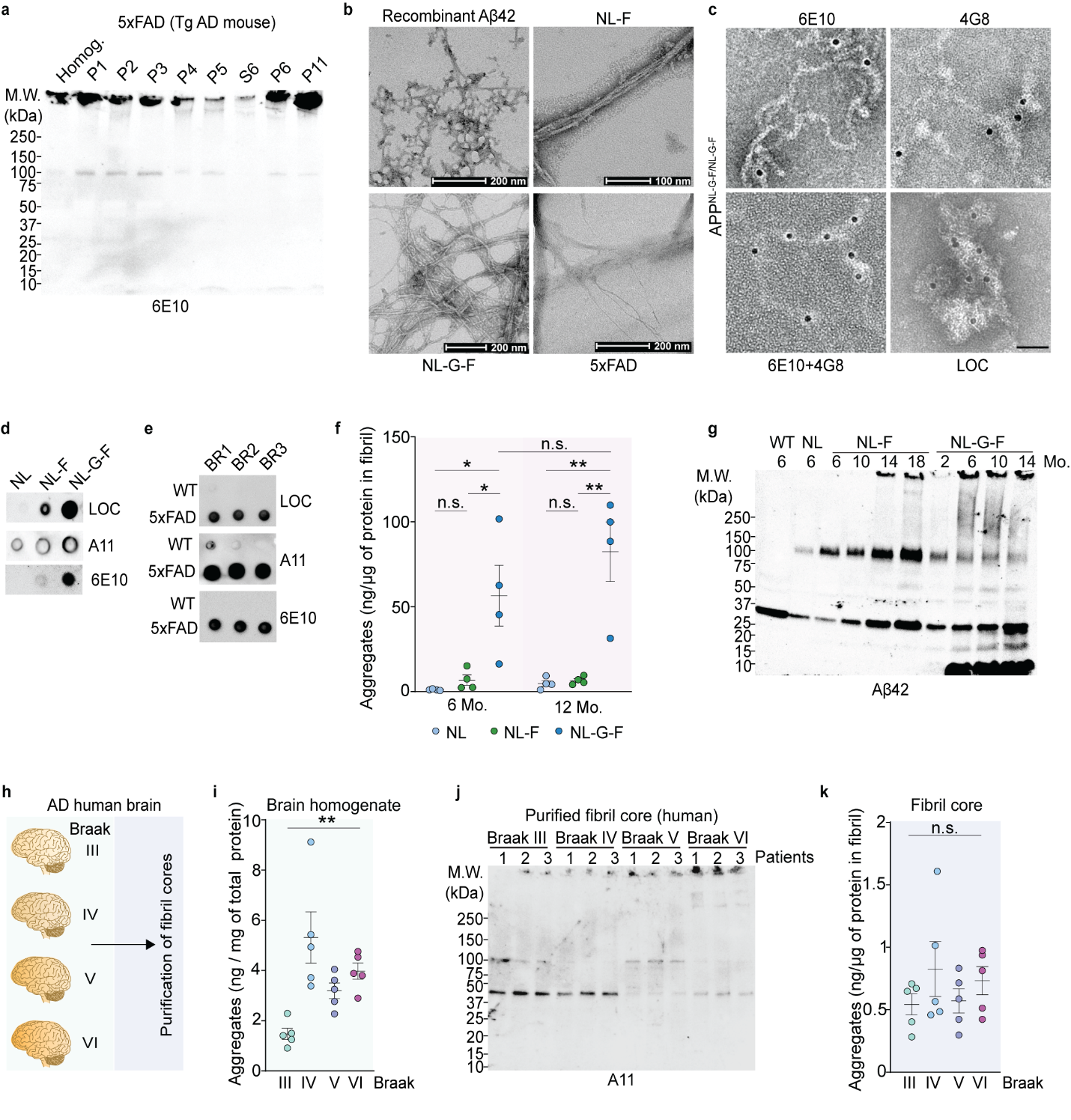
**Figure S1. Confirmation of amyloid fibril purification.**

(a) Representative WB across indicated fractions of transgenic 5xFAD cortical extracts collected during fibril purification using Aβ_1-16_ (6E10) antibody.

(b) Representative negative staining electron micrographs for *App^NL-F/NL-F^*, *App^NL-G-F/NL-G-F^*, and 5xFAD mouse brains and recombinant Aβ42 peptide fibrils as positive control.

(c) Panels of negative staining EM images showing immunogold labeling of NL-G-F fibrils using Aβ_1-16_ (6E10), Aβ_17-24_ (4G8), and anti-fibril (LOC) antibodies.

(d-e) Dot blot analysis of three *App KI* (*App^NL/NL^*, *App^NL-F/NL-F^*, and *App^NL-G-F/NL-G-F^*); and WT and transgenic 5xFAD brain amyloid fibrils using LOC, A11 and 6E10 antibodies.

(f) Sandwich ELISA-based quantification of Aβ aggregates in purified amyloid material isolated from the indicated cortical extracts from six- and twelve-month old *App* KI mouse brains.

(g) Representative WB indicating abundance of Aβ42 peptide monomers and higher order assemblies in purified amyloid extracted from all three *App KI* (*App^NL/NL^*, *App^NL-F/NL-F^*, and *App^NL-G-F/NL-G-F^*) brains and WT control at the indicated age.

(h) Summary of human amyloid purifications from postmortem AD brain cortex.

(i) ELISA-based quantification of Aβ aggregates in human brain cortex homogenates.

(j) WB analysis to probe A11-positive oligomeric assemblies in purified amyloid assemblies prepared from human brains.

(k) Quantification of total Aβ aggregates in human amyloid extracts using sandwich ELISA.

Data in f, i and k represents mean ± SEM; in f, n = four mice for each ELISA; in i and k n= five human brain samples analyzed. *, p-value < 0.05; **, p-value< 0.01; ***, p-value < 0.001; analyzed with unpaired Student’s t-test or one-way ANOVA with post-hoc Sidek test. BR = biological replicates; Tg = transgenic. P = pellet, S = supernatant. *App^NL/NL^* = NL, *App^NL-F/NL-F^* = NL-F, *App^NL-G-F/NL-G-F^* = NL-G-F. In c, scale bar: 50 nm

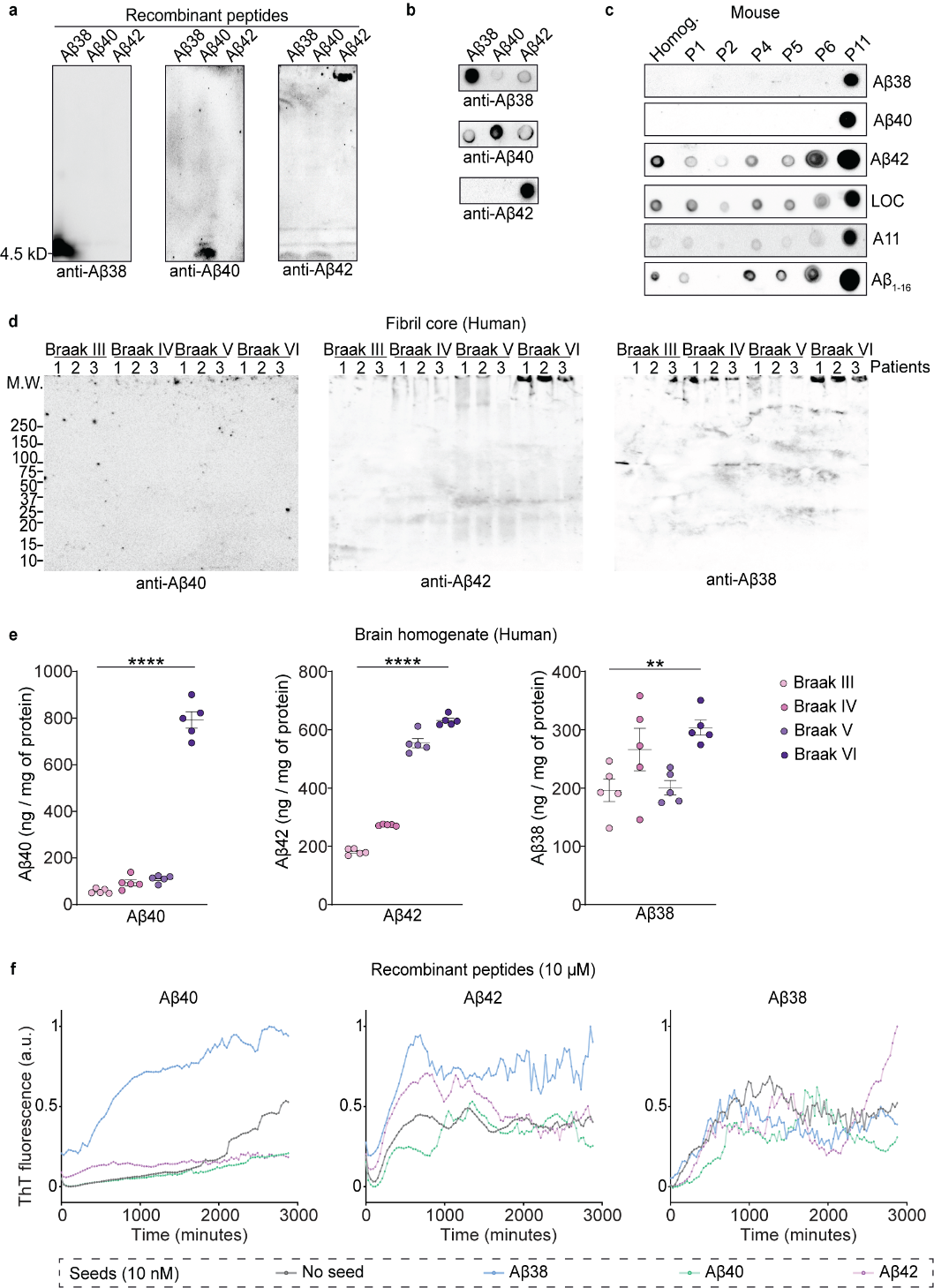

**Figure S2. Aβ38 peptides are present in high abundance in human and mouse fibrils**

(a-b) WB analysis and dot blots of recombinant Aβ38, Aβ40 and Aβ42 peptides using respective antibodies.

(c) Dot blot analysis of indicated fractions collected during amyloid fibril purification from *App^NL-G-F/NL-G-F^* mouse cortex using monomer-specific anti-Aβ38, Aβ40 and Aβ42 and conformation-specific antibodies.

(d) WB analysis of purified fibrils from human AD brains using Aβ38, Aβ40 and Aβ42 monomer-specific antibodies indicating the presence of Aβ38 and Aβ40 peptides.

(e) Quantification of Aβ38, Aβ40 and Aβ42 peptides in AD human cortical homogenates using sandwich ELISA.

(f) Kinetic plot of raw values of the ThT fluorescence intensities recorded in cross-seeding experiment. The curve was plotted similar to the those shown in Figure 3a-c, but without a secondary nucleation fit. Raw intensity values were normalized between zero and one before plotting the curve.

Data in e are mean ± SEM; n = five postmortem brain tissues for each group. *, p-value < 0.05; **, p-value< 0.01; ***, p-value < 0.001; analyzed with one-way ANOVA with post hoc Sidak test. P = pellet, and S = supernatant. *App^NL/NL^* = NL, *App^NL-F/NL-F^* = NL-F, *App^NL-G-F/NL-G-F^* = NL-G-F.

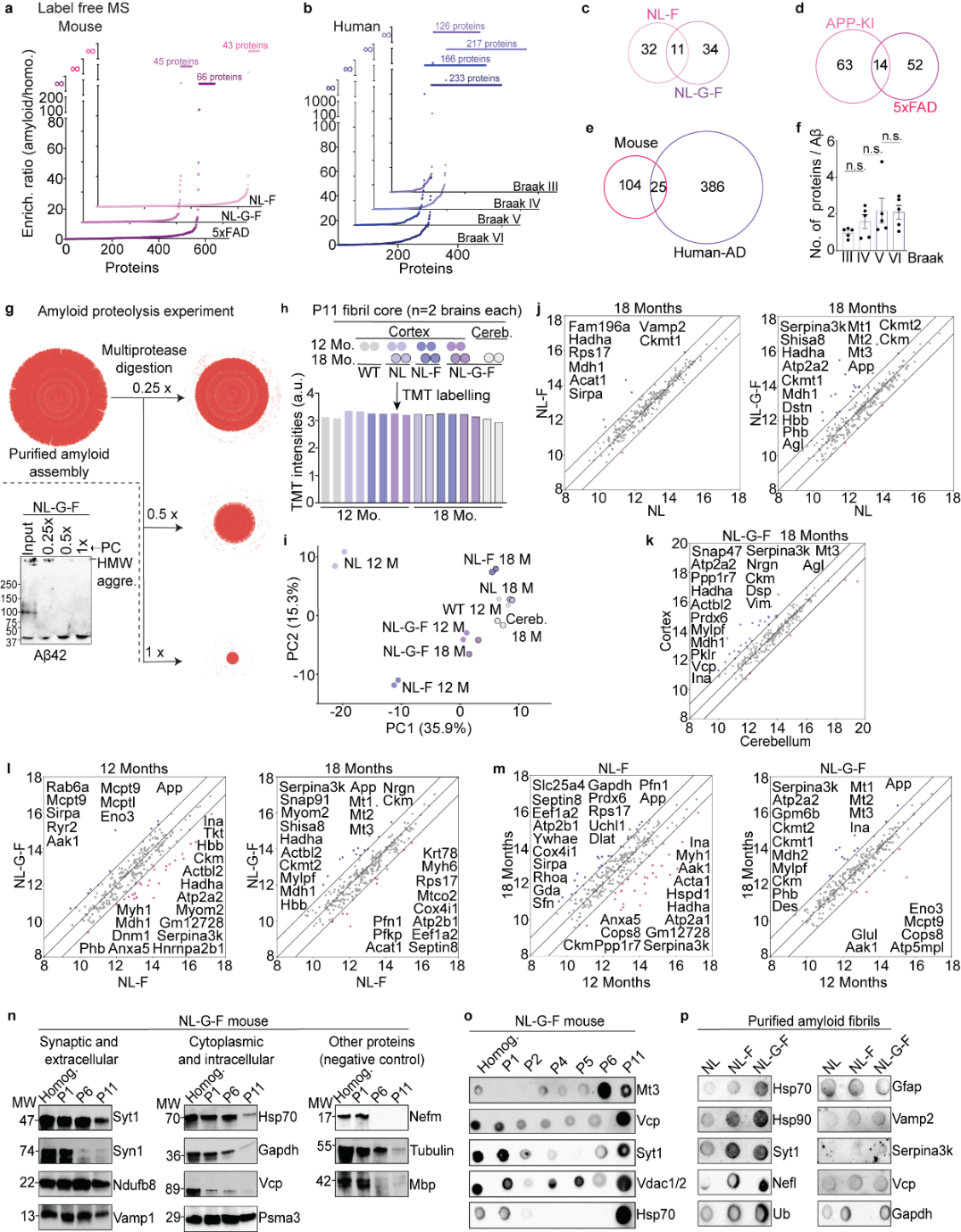

**Figure S3. Comprehensive MS analysis of purified mouse and human fibrils**

(a-b) Rank order plots of proteins for ratios between average NSAF values in purified fibrils compared to cortex homogenate samples in a label free MS analysis for indicated mouse and human fibril extracts.

(c-d) Venn diagram comparing proteins exclusively identified in purified amyloids prepared from indicated mouse strains.

(e) Venn diagram indicating number of proteins identified purified fibrils extracted from both human and mouse (*App^NL-F/NL-F^*, *App^NL-G-F/NL-G-F^*, 5xFAD) brain cortex.

(f) Number of proteins identified in human fibril samples compared to relative number of spec counts of APP peptides corresponding to Aβ region.

(g) Outline of multiprotease digestion experiment; representative blot in the inset showing concentration-dependent degradation of HMW amyloid assemblies following treatment with cocktail of multiple proteolytic enzymes: Arg-C, Asp-N, Lys-C, Glu-C, thermolysin, chymotrypsin, and trypsin.

(h) Experimental setup and relative TMT intensities across in 16-plex TMT channels. Each fibril source (mouse strain, age and brain region) has two biological replicates; each prepared from two mouse brains pooled together.

(i) PCA analysis of different biological samples using normalized TMT intensities of all identified proteins.

(j-m) Scatter plots comparing average TMT intensities of proteins measured in fibril cores extracted from twelve or eighteen months old *App KI* mouse cortical homogenates. A comparison of proteins identified in cortical and cerebellar fibrils is plotted in k.

(n-o) WBs and dot blot analysis of indicated fractions collected during amyloid fibril purification probing top protein interacting partners identified in MS analyses Few proteins (either not detected or non-specifically bound to fibrils) were immunoblotted as negative controls.

(p) Dot blots of relative levels of proteins in P11 fractions obtained from three *App KI* (*App^NL/NL^*, *App^NL-F/NL-F^*, *and App^NL-G-F/NL-G-F^*) brains.

P = pellet, and S = supernatant.

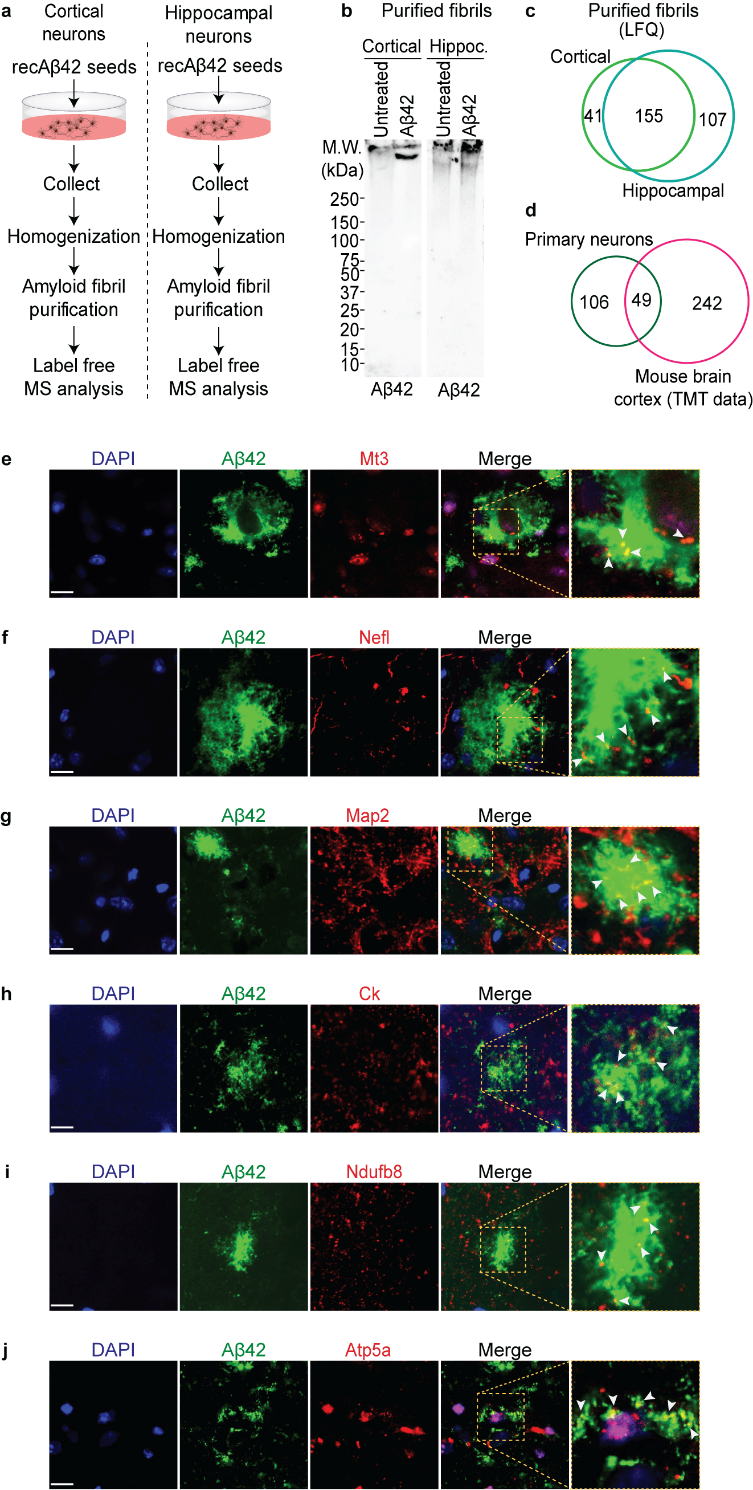

**Figure S4. *In vitro* and *in vivo* validation of proteomic data**

(a) Experimental outline of amyloid purification and supplementary MS analysis of fibrils collected from primary rat neurons seeded with recombinant Aβ42 peptides.
(b) Immunoblot confirmed formation of Aβ42-containing HMW aggregates in seeded neurons in comparison to unseeded neurons.

(c-d) Venn diagram comparing proteins identified in fibrils isolated from Aβ42 seeded hippocampal and cortical neurons; followed by proteins identified in neurons compared to those identified in fibrils purified from mouse cortex (TMT experiment dataset).

(e-j) Representative confocal images of IHC staining of three months old *App^NL-G-F/NL-G-F^* mouse brain cortex for most common proteins identified in multiple MS analysis confirming colocalization with large amyloid plaques stained with anti-Aβ42 antibody. Scale bar: 10 μm

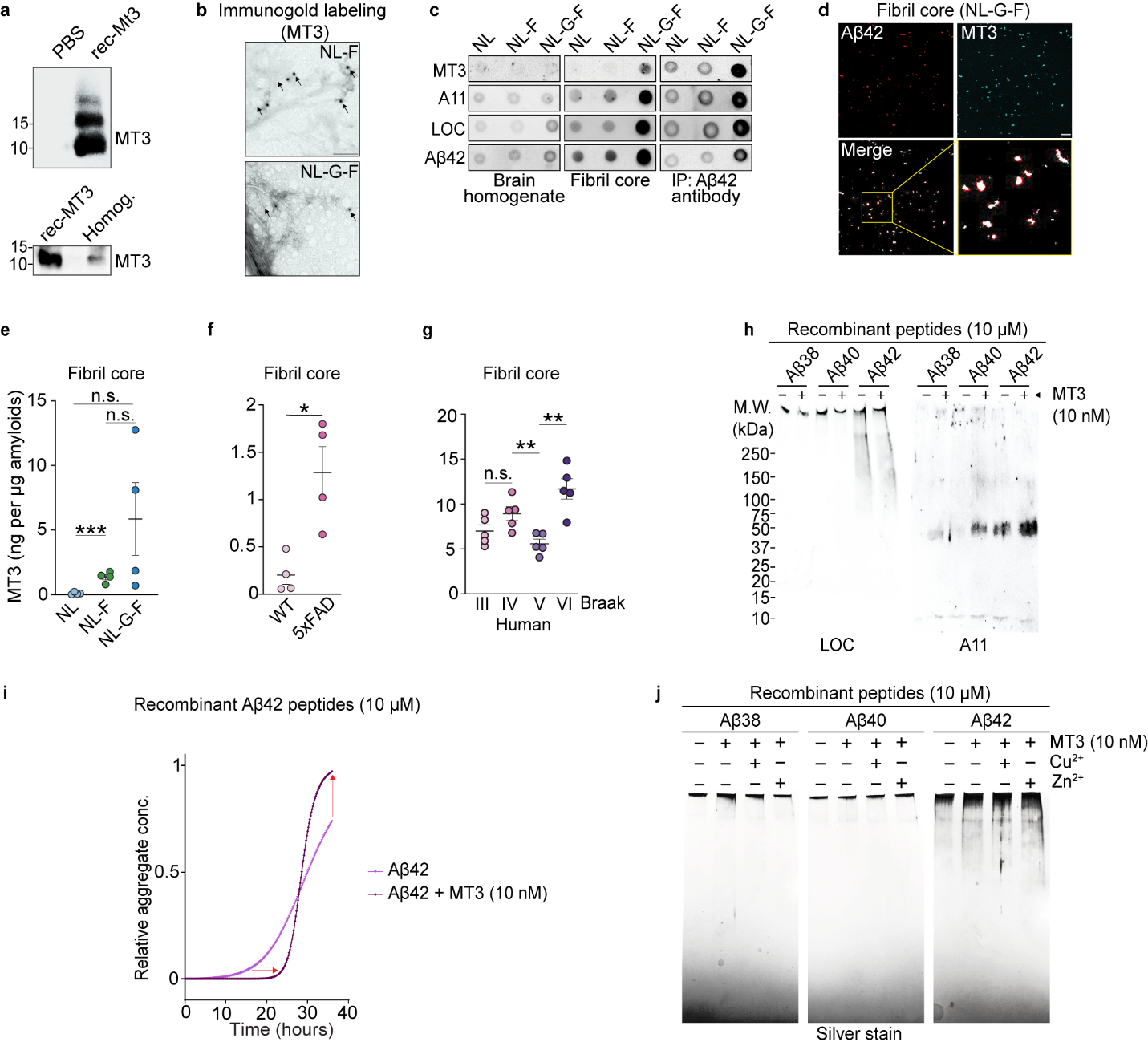

**Figure S5. Metallothionein-3, a MS identified protein affects amyloid aggregation**

(a) Confirmation blots to detect recombinant and mouse brain Mt3 protein using a monoclonal antibody.

(b) Immunogold labelling EM image panels confirming presence of Mt3 in purified amyloid fibrils isolated from *App^NL-F/NL-F^*, and *App^NL-G-F/NL-G-F^* brains.

(c) Dot blot analysis for relative abundance of the Mt3 protein in three *App KI* (*App^NL/NL^*, *App^NL-F/NL-F^*, and *App^NL-G-F/NL-G-F^*) brain homogenates and purified fibrils, with and without pull down using Aβ42 antibody.

(d) Immunostaining using monoclonal antibodies showing colocalization of Mt3 in Aβ42-positive purified fibrils.

(e-g) Sandwich ELISA for Mt3 protein in purified fibrils obtained from indicated mouse strains and human brain cortex homogenates.

(h) Immunoblots showing the effect of presence of Mt3 on aggregation of Aβ38, Aβ40 and Aβ42 peptides. All three peptides show increased oligomer formation (A11 blot). Presence of Mt3 shows least effect on inherent Aβ42 fibril formation compared to Aβ38, and Aβ40 peptides.

(i) ThT based *in vitro* aggregation kinetics experiment shows varying effects of 10 nM of Mt3 protein on the relative aggregation of 10 μM recombinant Aβ42 peptides.

(j) Silver stain image of SDS PAGE gels showing total amount of HMW aggregates of Aβ38, Aβ40, and Aβ42 formed under influence of Mt3, in absence of presence of copper (Cu^2+^) and zinc ((Zn^2+^) ions.

Data in e-g represents mean ± SEM; n = four for mouse and n= five for human patient sample. *, p-value < 0.05; **, p-value< 0.01; ***, p-value < 0.001; ****, p-value < 0.0001 analyzed with unpaired Student’s t-test. Scale bar: 10 μm

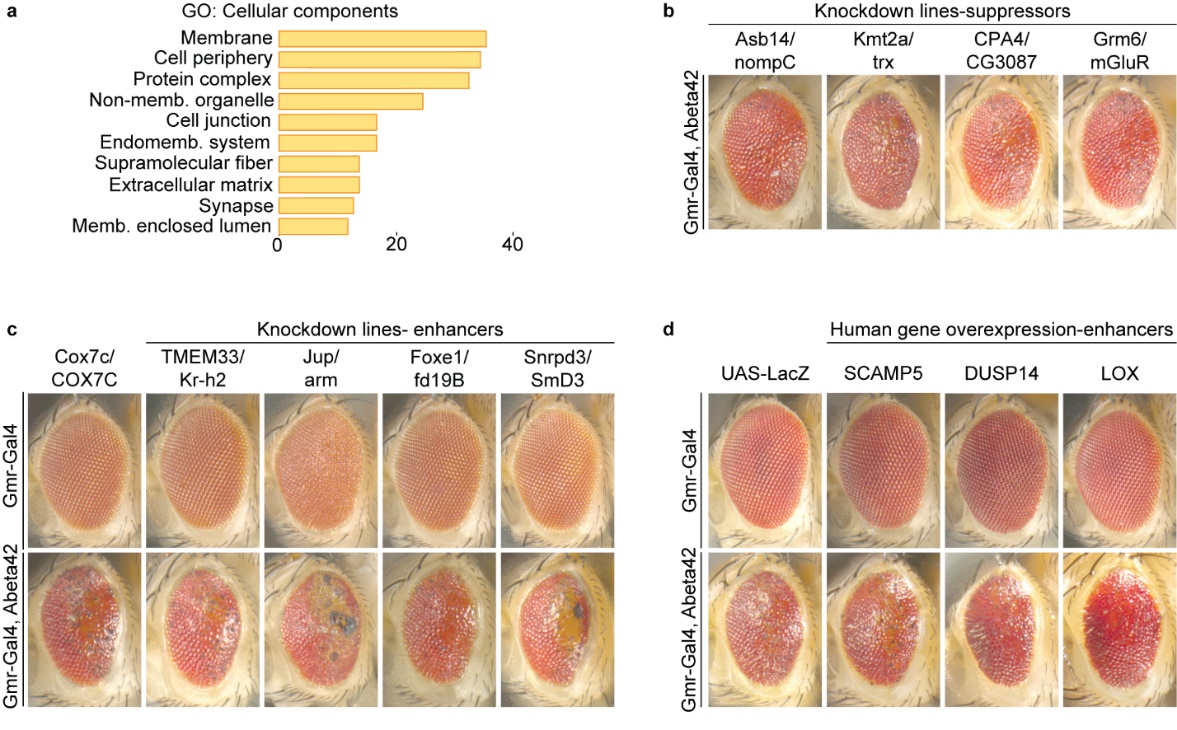

**Figure S6. Fly orthologues of MS-identified candidate proteins modulate Aβ42-induced neurotoxicity in vivo**

(a) GO enrichment analysis: cellular components for genes significantly enriched in Aβ42 flies compared to control flies showing abundance of membranous, extracellular, and synaptic proteins in amyloid fibrils.

(b) Representative eye images showing suppression of Aβ42-induced eye phenotypes upon expression of the indicated RNAi knockdown lines.

(c) Eye images showing enhancement of Aβ42-induced eye phenotypes upon expression of the indicated RNAi lines. Note that these lines have little to no effect on eye morphology in the absence of Aβ42 (top row).

(d) Overexpression of the indicated human transgenes enhances Aβ42-mediated phenotypes in the fly eye compared to the effect of the innocuous LacZ control transgene. These human genes do not disrupt the eye structures in the absence of Aβ42 (top row).

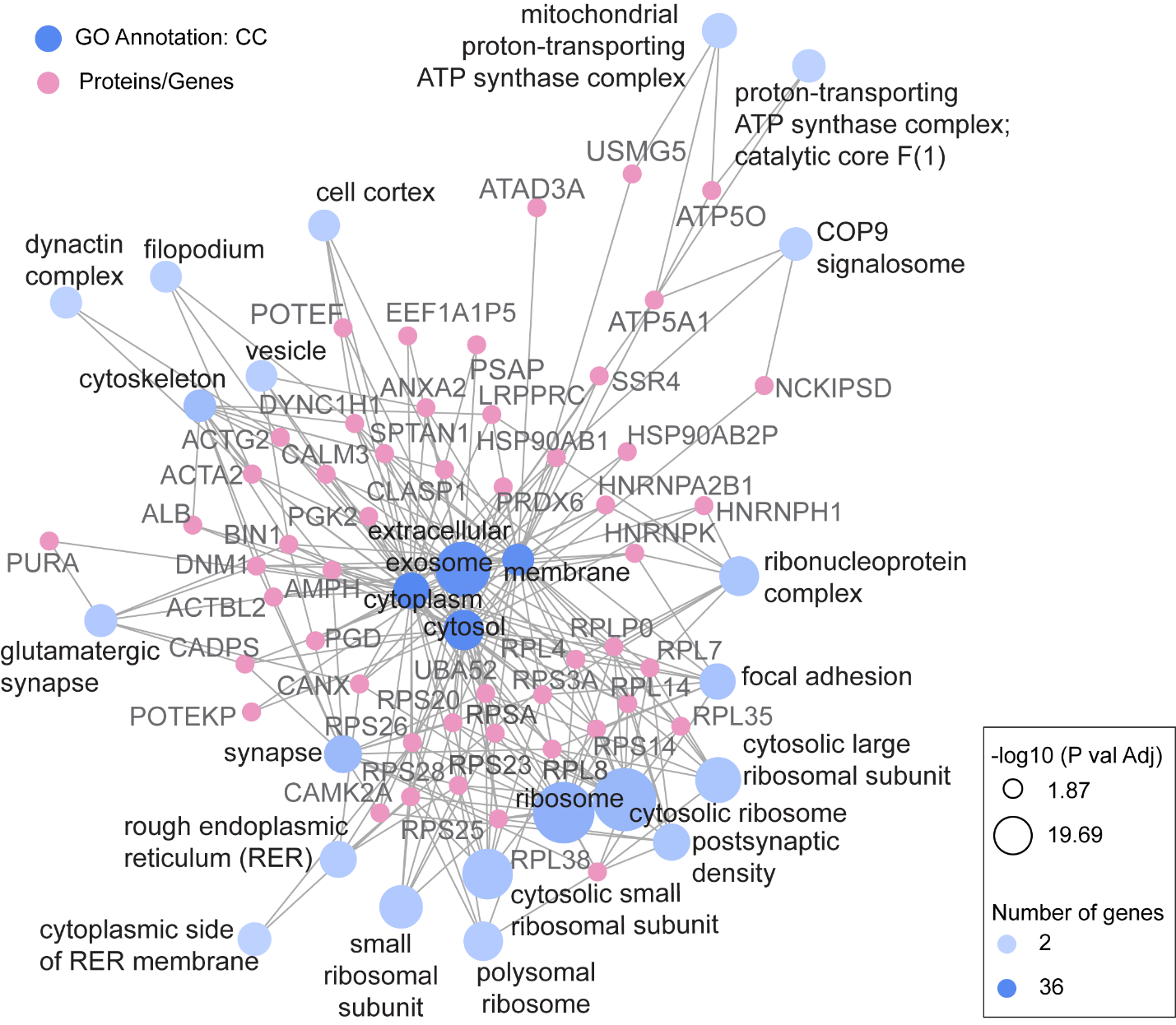

**Figure S7. Gene annotation based co-occurrence discovery**

Interactive protein interaction-based clusters network showing co-occurrence of proteins using GO: cellular component enrichment analysis of the consensus proteins (shown in Table 1 and 2). Top twenty-five GO annotations were chosen based on P-values and the network was drawn using online tool GeneCodis (<https://genecodis.genyo.es/>).

**Supplementary Tables**

**Table S1: List of proteins identified in label-free MS analysis of amyloid fibrils obtained from mouse cortices.** Proteins identified with a higher abundance in purified fibrils (n=four) obtained from *App^NL-F/NL-F^*, and *App^NL-G-F/NL-G-F^*, and 5xFAD mouse brains, 6 months age, compared to respective cortex homogenate as input (n=three). Average NSAF values for purified fibrils and cortex homogenates for individual proteins were taken into account for all calculations. Any protein not identified in purified fibrils samples dataset was not included in the table. Each sheet represents individual mouse line data sets, and each row has individual p values using Student’s t test, followed by adjusted p values with Benjamini-Hochberg (BH) correction. Number of significantly high abundance proteins in *App^NL-F/NL-F^*, and *App^NL-G-F/NL-G-F^*, and 5xFAD mouse are 56, 52, and 87, respectively. Experiment = specific data set, Uniprot accession= Uniprot identifier for each protein, ratio= log_2_ average NSAF values (purified fibrils/homogenate), t test p value = t test p value, Rank= rank ordered proteins based on p value (if p values are identical, higher ratio were given higher ranks), Adjusted p value is calculated using BH correction, description= protein description. Additional remarks indicate if results (increased abundance in purified fibrils) are statistically significant or not.

**Table S2: List of proteins identified in label free MS analysis of amyloid fibrils obtained from human cortices.** Proteins identified with a higher abundance in purified fibrils obtained from human brain cortices, AD Braak stages III to VI, five patients each, compared to respective cortex homogenate as input (n=five). Average NSAF values for purified fibrils and cortex homogenates for individual proteins were considered for comparisons. Proteins not identified in purified fibril samples dataset were not included in the table. Each sheet represents individual mouse line data sets, and each row has individual p values using Student’s t test, followed by adjusted p values with BH correction. Number of significantly high abundance proteins in Braak stage III, IV, V, and VI are 84, 142, 115 and 148, respectively. Experiment = specific data set, Uniprot accession= Uniprot identifier for each protein, ratio= log_2_ average NSAF values (purified fibrils/homogenate), t test p value = t test p value, Rank= rank ordered proteins based on p value (if p values are identical, higher ratio were given higher ranks), Adjusted p value is calculated using BH correction, description= protein description. Additional remarks indicate if results (increased abundance in purified fibrils) are statistically significant or not.

**Table S3: List of proteins identified in label-free MS analysis of amyloid fibrils obtained from mouse and human cortices following multiprotease digestion.** Proteins identified with a higher abundance in purified fibrils following their additional multiprotease digestion (n=4) compared to those without this additional lysis step (n=4 each). Average NSAF values of individual proteins were considered for comparisons between both sample types. Proteins only identified in multiprotease digestion sample datasets were included in the table. Each sheet represents individual mouse line data sets, and each row has individual p values using Student’s t test, followed by adjusted p values with BH correction. Number of significantly high abundance proteins in NL, NL-F, NL-G-F, and 5xFAD mouse are 5, 9, 15, and 89 while in human samples the numbers are 42, 151, 105, and 352 for Braak stage III, IV, V, and VI, respectively. Experiment = specific data set, Uniprot accession= Uniprot identifier for each protein, ratio= log_2_ average NSAF values (purified fibrils/homogenate), t test p value = t test p value, Rank= rank ordered proteins based on p value (if p values are identical, higher ratio were given higher ranks), Adjusted p value is calculated using BH correction, description= protein description.

**Table S4: List of proteins identified in multiplex TMT analysis of amyloid fibrils obtained from mouse of different age groups.** Proteins identified in 16-plex TMT analysis containing eight biological conditions, each in two biological replicates. Average normalized TMT intensity values have been used for making comparisons between individual conditions. No intensity cutoff is applied. Each sheet shows comparisons between individual biological groups and proteins only in higher abundance in every comparison shown in the table. Experiment = specific data set, Uniprot accession= Uniprot identifier for each protein, ratio= log_2_ average TMT intensity (group 1/group 2), protein= protein name, description= protein description.

**Table S5: List of proteins identified in label-free MS analysis of amyloid fibrils obtained from rat hippocampal and cortical neurons incubation recombinant Aβ42 seeds.** Proteins identified in fibrils purified independently from rat cortical and hippocampal neurons (n= four). Proteins identified at least in two independent fibril preparation from each culture type have been considered and the table shows only those proteins that were also identified in TMT analysis (shown in table S4). Experiment = specific data set, Uniprot accession= Uniprot identifier for each protein, occurrence score= number of occurrences in independent fibril preparations, description= protein description.

**Table S6: List of proteins identified in label-free MS analysis of amyloid fibrils obtained from Aβ42 flies.** Proteins identified with a higher abundance in purified fibrils obtained from Aβ42 flies, compared to control Lac-Z flies (n = four for both). In sheet 1, average NSAF values for purified fibrils for individual proteins across two groups were considered for comparisons, each row has individual p values using Student’s t test, followed by adjusted p values with BH correction. Corresponding human and mouse orthologs have been identified for each fly gene (https://www.flyrnai.org/diopt). Proteins with significantly high abundance in Aβ42 flies are shown in the table. Sheet 2 shows proteins considered for obtaining RNAi lines following identification of Fly orthologs based on scoring. Experiment = specific data set, Uniprot accession= Uniprot identifier for each protein, ratio= log_2_ average NSAF values of purified fibrils (Aβ42 /control flies), protein= protein names, t test p value = t test p value, Rank= rank ordered proteins based on p value (if p values are identical, higher ratio were given higher ranks), Adjusted p value is calculated using BH correction, description= protein description, putative human ortholog= human gene orthologous to the identified fly genes, putative mouse ortholog= mouse gene orthologous to the identified fly genes obtained using Dipot online tool, only high and moderate scoring genes were considered in the analysis). Additional remarks indicate if results (increased abundance in purified fibrils) are statistically significant or not.
