## Additional File 2 for "Amyloid fibril proteomics of AD brains reveals modifiers of aggregation and toxicity"

**SUPPLEMENTARY INFORMATION FILE 2**

**Supplementary Materials and Methods**

**Human samples**

Additional details on AD patients: their diagnosis, and neuropathological conditions are provided in Supplementary methods:

| **S.N.** | **Age** | **Sex** | **PMT (hours)** | **A** | **B** | **C** | **Braak stage** | **Neuropathological condition** |
| --- | --- | --- | --- | --- | --- | --- | --- | --- |
| 1 | 91 | F | 8 | 2 | 2 | 1 | 3 | AD neuropathologic change, intermed |
| 2 | 93 | M | 25 | 3 | 2 | 1 | 3 | AD neuropathologic change, intermed |
| 3 | 92 | M | 4 | 3 | 2 | 2 | 3 | AD neuropathologic change, intermed |
| 4 | 100 | F | 13 | 2 | 2 | 1 | 3 | AD neuropathologic change, intermed |
| 5 | 90 | M | 6 | 2 | 2 | 2 | 3 | AD neuropathologic change, intermed |
| 6 | 93 | M | 13 | 3 | 2 | 2 | 4 | AD neuropathologic change, intermed |
| 7 | 68 | F | 0 | 3 | 2 | 3 | 4 | AD neuropathologic change, intermed |
| 8 | 91 | F | 13 | 3 | 2 | 2 | 4 | AD neuropathologic change, intermed |
| 9 | 102 | F | 6.5 | 3 | 2 | 3 | 4 | AD neuropathologic change, intermed |
| 10 | 88 | F | 5 | 2 | 2 | 2 | 4 | AD neuropathologic change, intermed |
| 11 | 84 | M | 8 | n.a. | 3 | 3 | 5 | AD neuropathologic change, high |
| 12 | 93 | M | 5.5 | 3 | 3 | 3 | 5 | AD neuropathologic change, high |
| 13 | 85 | F | 10 | 3 | 3 | 3 | 5 | AD neuropathologic change, high |
| 14 | 81 | F | 7 | 3 | 3 | 3 | 5 | AD neuropathologic change, high |
| 15 | 88 | M | 10.5 | 3 | 3 | 3 | 5 | AD neuropathologic change, high |
| 16 | 85 | M | 14 | 3 | 3 | 3 | 6 | AD neuropathologic change, high |
| 17 | 87 | M | 6.5 | 3 | 3 | 3 | 6 | AD neuropathologic change, high |
| 18 | 82 | F | 5 | 3 | 3 | 3 | 6 | AD neuropathologic change, high |
| 19 | 82 | F | 4 | 3 | 3 | 3 | 6 | AD neuropathologic change, high |
| 20 | 63 | M | 18.5 | 3 | 3 | 3 | 6 | AD neuropathologic change, high |

**Antibodies**

Antibodies used: antifibril LOC antibody: EMD Millipore #AB2287, RRID:AB_11211951; oligomer A11 antibody: Thermo Fisher Scientific #AHB0052, RRID:AB_2536236; Aβ_1-16_ (6E10): Biolegend #803001, RRID:AB_2564653; Aβ_17-24_ (4G8): Biolegend #800701, RRID:AB_2564633; Aβ_1-38_: Biolegend #943302, RRID:AB_2890865; Aβ1-40: GeneTex #GTX40068, RRID:AB_423909; Aβ1-42: Santa Cruz Biotecnology #sc-28365, RRID:AB_626669; APP: Thermo Fisher Scientific #OMA1-031232, RRID:AB_ 325526; MT3: Thermo Scientific #PA5-50448, RRID:AB_2635901; creatine kinase: Abcam # ab174672, RRID:AB_2747709; Atp5a: Abcam # ab14748, RRID:AB_301447; actin: Santa Cruz Biotechnology #sc-8432, RRID:AB_626630; Syt1: Synaptic Systems #105-003, RRID:AB_2619763; Syn1: Millipore Sigma #AB1543, RRID:AB_2200400; Ndufb8: Abcam #ab192878, RRID:AB_2847808; Vamp2: Synaptic systems #104 202, RRID: AB_887810; Hsc70: Santa Cruz Biotecnology #sc-7298, RRID:AB_627761; Millipore Sigma #Hsp70: SAB4200714; Hsp90: Cell Signaling Technology #4874S, RRID:AB_2121214; GAPDH: Santa Cruz Biotecnology #sc-47724, RRID:AB_627678; VCP: Abcam #ab111740, RRID:AB_10861709; Psma3: Cell Signaling Technology #2456S, RRID:AB_2171417; Nefm: Cell Signaling Technology #2838T, RRID:AB_561191; Nefl: Abclonal #A0257, RRID:AB_2757070; β-tubulin: Novus Biologicals #NB100-1612, RRID:AB_10000548; Mbp: Novus Biologicals NB600-717, RRID:AB_531559; Vdac1/2: Abcam #ab34726, RRID:AB_778788; Ubiquitin: Santa Cruz Biotecnology #sc-8017, RRID:AB_628423; Serpina3k: Proteintech #55480-1-AP, RRID:AB_2881346; Gfap: Novus Biologicals #NBP1-05198, RRID:AB_1556315; Map2: Millipore Sigma #AB5543, RRID:AB_571049

**MS data acquisition**

We acquired MS data following previously described methods and parameter settings [1, 2]. Dried peptide samples were resuspended in 20 µl peptide resuspension buffer (94.875% H2O with 5% ACN and 0.125% FA) and 3 µg of each sample as determined by micro BCA assay (Thermo Fisher Scientific, Cat#23235), was loaded via autosampler with either Thermo EASY nLC 100 UPLC or UltiMate 3000 HPLC pump, onto a vented Pepmap 100, 75 µm × 2 cm, nanoViper trap column coupled to a nanoViper analytical column (Thermo Fisher Scientific) with a stainless steel emitter tip assembled on the Nanospray Flex Ion Source with a spray voltage of 2,000 V. MS data was acquired via a coupled Orbitrap Fusion mass spectrometer with combination of a quadrupole, an Orbitrap and a linear ion trap mass spectrometer ("tribrid"). For label-free quantification, following MS parameters were used: ion transfer tube temp, 300°C; Easy-IC internal mass calibration; default charge state, 2; cycle time, 3 s; detector type set to Orbitrap; 60K resolution; wide quad isolation; mass range, normal; scan range, 300–1,500 m/z; maximum injection time, 50 ms; AGC (automatic gain control) target, 200,000; microscans, 1; S-lens RF level, 60; without source fragmentation; and datatype, positive and centroid; MIPS set as on; included charge states, 2–6 (reject unassigned); dynamic exclusion enabled, with *n* = 1 for 30- and 45-s exclusion duration at 10 ppm for high and low; precursor selection decision, most intense, top 20; isolation window, 1.6; scan range, auto normal; first mass, 110; and collision energy, 30%. For collision-induced dissociation (CID): detector type, ion trap; OT resolution, 30K; IT scan rate, rapid; maximum injection time, 75 ms; AGC target, 10,000; Q, 0.25; and inject ions for all available parallelizable time. For absolute quantification of Aβ42 peptides, each sample was run one more time using the targeted MS/MS approach by providing a list of selected m/z values, corresponding to Aβ42 peptides. This approach circumvents the problem often associated with data-dependent exclusion of highly abundant peptides (like Aβ peptides in fibrils) and can simultaneously quantify the Aβ peptides with high resolution.

For TMT samples, each fraction of multiplexed peptides was resuspended in 20 μL of peptide resuspension buffer, and following micro-BCA quantification, three micrograms of each fraction were analyzed separately on tribrid Orbitrap Fusion MS system. The chromatographic run was performed with a 4 h gradient beginning with 100% buffer A and 0% B and increased to 7% B over 5 min, then to 25% B over 160 min, 36% B over 40 min, 45% B over 10 min, 95% B over 10 min, and held at 95% B for 15 min before terminating the scan. Buffer A contained 5% acetonitrile (ACN) and 0.125% formic acid in H2O, and buffer B contained 99.875 ACN with 0.125% formic acid. To enhance to accuracy and sensitivity of MS analysis, a multi-notch MS3 method was applied with the following parameters: ion transfer tube temp = 300 °C, easy-IC internal mass calibration, default charge state = 2, and cycle time = 3 s. MS1 detector was set to orbitrap with 60 K resolution, wide quad isolation, mass range = normal, scan range = 300–1800 m/z, max injection time = 50 ms, AGC target = 6 × 105, microscans = 1, RF lens = 60%, without source fragmentation, and datatype = positive and centroid. Monoisotopic precursor selection was set to include charge states 2–7 and reject unassigned. Dynamic exclusion was allowed; n = 1 exclusion for 60 s with 10 ppm tolerance for high and low. The intensity threshold was set to 5 × 103. Precursor selection decision = most intense, top speed, 3 sec. MS2 settings include isolation window = 0.7, scan range = auto normal, collision energy = 35% CID, scan rate = turbo, max injection time = 50 ms, AGC target = 6 × 105, and Q = 0.25. In MS3, the top 10 precursor peptides selected for analysis were then fragmented using 65% higher-energy collisional dissociation before orbitrap detection. A precursor selection range of 400–1200 m/z was chosen with mass range tolerance. An exclusion mass width was set to 18 ppm on the low and 5 ppm on the high. Isobaric tag loss exclusion was set to TMT reagent. Additional MS3 settings include an isolation window = 2, orbitrap resolution = 60 K, scan range = 120–500 m/z, AGC target = 6 × 105, max injection time = 120 ms, microscans = 1, and datatype = profile.

**MS data analysis**

For label-free quantification, each MS raw file was extracted using the offline Rawconverter tool ([http://fields.scripps.edu/rawconv](http://fields.scripps.edu/rawconv/)/) to obtain MS1 and MS2 files. These files were used as input for identification, quantification, and detailed analyses of the data on Integrated Proteomics Pipeline - IP2 (Bruker: <http://www.integratedproteomics.com/>), a web-based MS data search engine. We downloaded the mouse, human, rat, or fly proteome database from Uniprot. For the *App KI* mice, additional App sequences with the specific mutations for NL, NLF, and NL-G-F mice were added prior to the identification of peptides using ProLuCId and SEQUEST algorithms. We used peptide mass tolerances of 20 ppm of precursor and 600 ppm for fragment ions. Fully and half-tryptic peptide candidates were included in the search space, all that fell within the mass tolerance window with no miscleavage constraint, assembled, and filtered with DTASelect2 (ver. 2.1.3). Static modifications at 57.02146 C were included. The target-decoy strategy was used to verify peptide probabilities and false discovery ratios. A minimum peptide length of five was set for the process of each protein identification, and each dataset included a 1% FDR rate at the protein level based on the target-decoy strategy.

Data analysis for TMT-MS was performed as previously described [1]. In summary, protein identification, TMT quantification, and analysis were performed with The Integrated Proteomics Pipeline-IP2 (Integrated Proteomics Applications, Inc., <http://www.integratedproteomics.com/>). Proteomic results were analyzed with ProLuCID, DTASelect2, Census, and QuantCompare. MS1, MS2, and MS3 spectrum raw files were extracted using RawExtract 1.9.9 software (<http://fields.scripps.edu/downloads.php>). Pooled spectral files from all eight fractions were searched against the Uniprot mouse protein database and matched to sequences using the ProLuCID/SEQUEST algorithm (ProLuCID ver. 3.1) with 50 ppm peptide mass tolerance for precursor ions and 600 ppm for fragment ions. Other parameters were similar to LFQ analysis except for additional static modification 304.2071 K for TMTpro were included. The target-decoy strategy was used to verify peptide probabilities and false discovery ratios. Isobaric labeling analysis was established with Census 2 as previously described [3] Normalized TMT intensities of individual proteins identified across sixteen channels in the analysis were used for PCA analysis in the Perseus data analysis tool, while the means of the normalized TMT intensities for each experimental group were used for further data analysis.

[Gene](https://www.sciencedirect.com/topics/biochemistry-genetics-and-molecular-biology/gene-ontology) enrichment Ontology (GO) based statistical overrepresentation analyses were performed using ShinyGO 0.76.3 (<http://bioinformatics.sdstate.edu/go/>) online tool [4]. Specifically, for the label-free experiment, we enlisted proteins identified exclusively in purified amyloid fibrils (the query) and used those to identify overrepresented for cellular components against a reference list comprising aggregated total proteins identified in LC-MS/MS analysis of respective inputs. A similar analysis for grouping proteins significantly higher in fibrils purified from Aβ42 flies compared to negative controls was performed. Protein ontologies with Fisher statistical tests with false discovery rate correction less than 0.05 were considered significant in these analyses.
